# The HUSH Complex Dictates EBV-transformed B cell Sensitivity to NK Cell Surveillance Through Repression of NKG2A Ligand γ-Proto-cadherin

**DOI:** 10.64898/2026.06.10.731186

**Authors:** Yizhe Sun, Isabella Y Kong, Wanyu Li, Jesse S. Plung, Eric M Burton, Ling Zhong, Shunji Li, Herman van Besien, Suhong Sun, Hugh T. Reyburn, Jonathan Abraham, Lisa Giulino-Roth, Benjamin E. Gewurz

## Abstract

Natural Killer (NK) cells control Epstein-Barr virus (EBV), though how EBV+ B-cells escape NK surveillance to form tumors remains unknown. To gain insights, we performed a human genome-wide CRISPR-Cas9 screen in EBV-transformed lymphoblastoid cell lines (LCL). This revealed that the HUSH complex maintains LCL sensitivity to NK. HUSH knockout (KO) de-repressed LCL protocadherin gamma (PCDHG), typically expressed by neurons. PCDHG expression protected LCLs from NK and was necessary for HUSH KO-driven NK resistance.

CRISPR analyses revealed NKG2A/CD94 as the PCDHG NK inhibitory counter-receptor, and NKG2A KO restored NK lysis of HUSH KO LCLs. HUSH perturbation impaired NK control of murine LCL xenografts *in vivo.* A subset of follicular lymphoma (FL) upregulate PCDHG, and PCDHG KO enhanced FL lysis by NK. Therefore, HUSH plays dual roles in foreign DNA surveillance and in support of NK, potentially as a neuronal ‘don’t kill me’ signal that can be exploited by transformed cells.

## Introduction

Epstein-Barr virus (EBV) persistently infects >95% of adults worldwide, causes infectious mononucleosis (IM) and is associated with over 200,000 cancer cases annually^1–4^. These include Burkitt and Hodgkin lymphomas, acquired immunodeficiency syndrome (AIDS)-associated B cell lymphomas and post-transplant lymphoproliferative disease (PTLD)^5^.

EBV-associated lymphomas occur most frequently in individuals with primary or acquired immunodeficiency, highlighting cellular immunity as playing a key role in control of acute and chronic EBV infection^5–10^.

Natural killer (NK) cells bridge innate and adaptive immune responses and are essential for immune surveillance and control of EBV infection^11–13^. Primary NK cell deficiencies can manifest by fulminant EBV infection, elevated EBV viral loads and the development of EBV-driven B cell lymphomas^11,14–16^. NK cells retain protective function even with T-cell immunodeficiency, including following stem cell transplantation, where NK cells reconstitute in advance of T cells^17,18^.

In the absence of cell-mediated immune control, EBV transforms primary human B cells into immortalized, continuously proliferating lymphoblastoid cell lines (LCLs)^19^. LCLs express all eight EBV latency oncoproteins, which upregulate a range of activating receptors and render LCLs highly sensitive to immune surveillance^20,21^. For example, CD48 is one of the first and most highly EBV-upregulated plasma membrane proteins in newly infected B cells^22–24^ and interacts with the activating receptor 2B4 on T and NK cells. Although Epstein-Barr nuclear antigen 1 (EBNA1) suppresses NK activation early after infection by downregulating several NK activating ligands^25^, this interaction is critical for immune control of EBV infection as genetic deficiency in SLAM-associated protein (SAP), the signaling adapter protein used by 2B4 leads to fulminant EBV-driven mononucleosis, hemophagocytic lymphohistiocytosis, hypogammaglobulinemia, and B-cell lymphoma.

PTLD is a major complication of 1-20% of solid and allogeneic bone marrow transplants^26,27^.

Early PTLD is characterized by EBV-driven polyclonal B-cell proliferation without evidence of malignant transformation^28^. Reduction of immunosuppression in patients with early lesions often leads to significant clinical improvement^29^. However, with the accumulation of additional genetic mutations, PTLD can evolve into monomorphic, clonal lymphoma^28^. Although reducing immunosuppression is first-line therapy, this often does not cause durable remission^30^, suggesting that tumor immune escape mechanisms are active in these cases.

NK cell immune surveillance depends on the balance of signaling from activating versus inhibitory receptors, which interact with the repertoires of cognate ligands expressed by different target cells. The major activating receptors expressed by human NK cells include NKG2D, the NCRs, 2B4 and DNAM-1, while the major inhibitory receptors are the KIR and CD94/NKG2A receptors that bind classical MHC Class I and the non-classical MHC class I molecule HLA-E respectively^31^. When activating signals outweigh inhibitory signals, NK cells become activated and initiate the killing process^32–36^. As mentioned previously, the 2B4 receptor is critical for NK recognition of EBV-transformed B-cells, but little is known as to whether EBV can modulate inhibitory ligand expression to evade NK surveillance. We have undertaken a human genome-wide CRISPR-Cas9 loss-of-function screen to define resistance pathways that allow EBV-transformed B cells to evade NK attack. In addition to CD48 and LFA-1, all components of the HUSH epigenetic silencing complex scored as top screen hits. Perturb-seq highlighted that HUSH knockout (KO) upregulated expression of multiple proto-cadherin gamma proteins (PCDHGs), which were required for HUSH KO LCL protection from NK. A CRISPR screen in NK cells identified NKG2A (also called KLRC1 or CD159a) as the NK-cell inhibitory receptor, which together with KLRD1/CD94 physically associated with PCDHG.

## Results

### CRISPR/Cas9 screen reveals key LCL pathways that support NK surveillance

We confirmed that LCLs are highly sensitive to lysis by NK cells. At an effector to target ratio of 1:1, the NK-cell line YT-INDY^37^ or interleukin-15/nicotinamide (NAM) activated primary human NK cells^38^ both killed the majority of GM15892 LCLs after 5 days of co-culture (**Fig. S1A**). To identify LCL genes that regulate NK cell cytotoxicity, we performed a human genome-wide CRISPR-Cas9 loss-of-function screen (**Fig. 1A-B**). Cas9+ GM15892 LCLs were transduced with the Brunello lentivirus single-guide RNA (sgRNA) library^39^, comprised of 77,441 sgRNAs, at a multiplicity of infection of 0.3. Most human genes are targeted by four independent Brunello sgRNAs.

**Figure 1.**
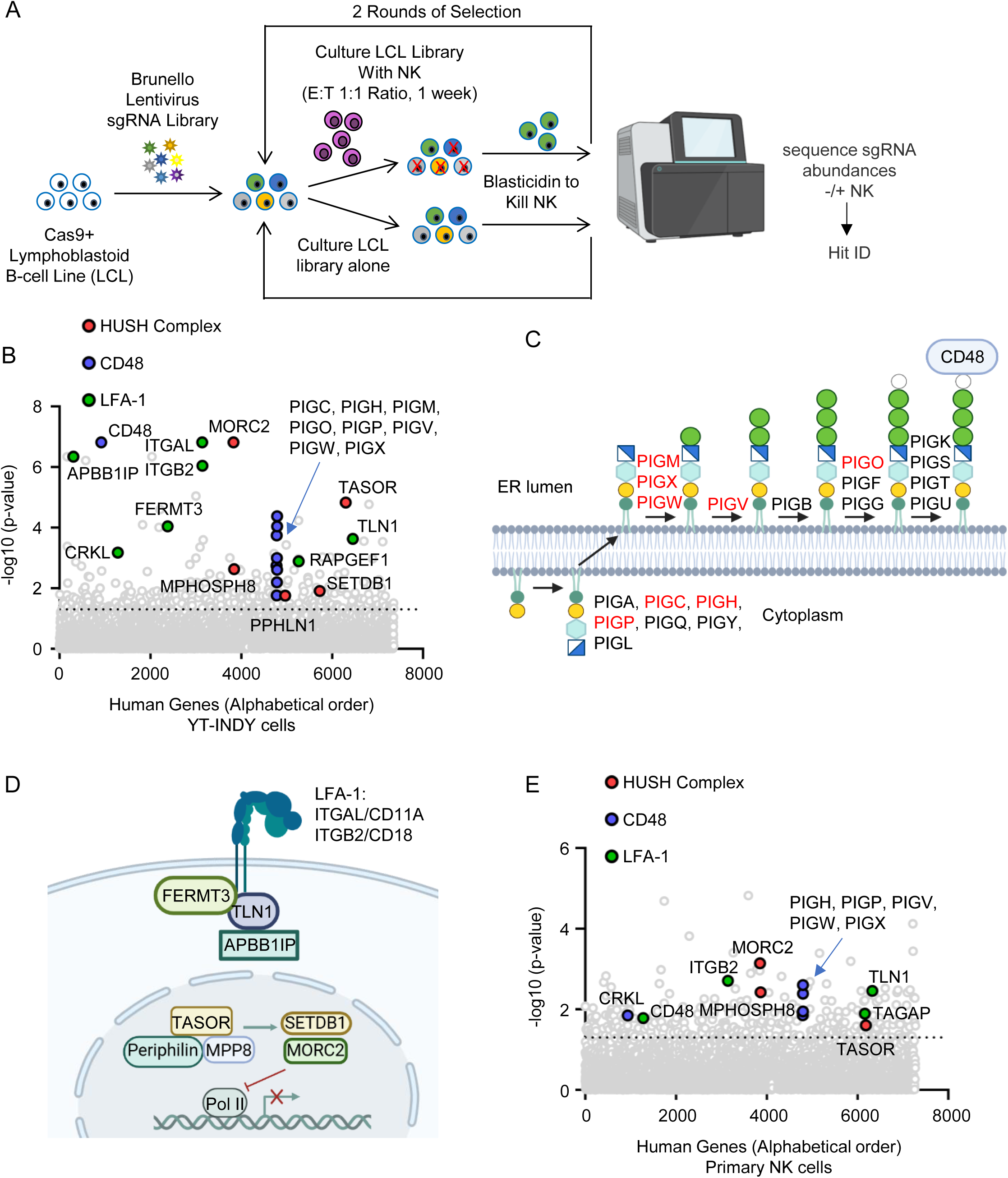
Human Genome-wide CRISPR/Cas9 Screen for LCL Intrinsic Factors that Support NK Cytotoxicity. A) CRISPR screen workflow. Cas9+ GM15892 cells were transduced with the Brunello sgRNA library and incubated with NK cells for 7 days. NK cells were then removed by 7 days treatment of blasticidin (Cas9+ GM15892 are resistant). After two rounds of selection, genomic DNA was harvested from surviving GM15892 versus from GM15892 cultured in parallel without NK and used for sgRNA quantitation. B) YT-INDY NK screen volcano plot analysis, using data from three independent CRISPR screen replicates. Selected screen hit categories are highlighted. Dotted line, adjusted p-value=0.05. C) Schematic highlight GPI anchor biosynthesis pathway genes that scored in the screen (red). CD48 is a GPI-anchored plasma membrane protein. D) Schematic depicting the two groups of enriched hits: (1) LFA-1 related genes and (2) HUSH complex related genes. E) IL-15/NAM treated primary human NK cell screen volcano plot analysis, using data from two independent CRISPR screen replicates. Selected screen hit categories are highlighted. Dotted line, adjusted p-value=0.05.

Successfully transduced cells were puromycin selected and cultured alone or co-cultured with YT-INDY for 7 days at an effector:target ratio of 1:1. Cultures were treated with blasticidin for 7 days to remove the NK cells (Cas9+ GM15892 are blasticidin resistant). Surviving cultures were cultured alone or with fresh NK cells for two more rounds (**Fig. 1A**). Surviving LCLs from two screen replicates were harvested, and the abundance of PCR-amplified sgRNAs were quantitated by next-generation sequencing.

We used the STARS algorithm^40^ to identify statistically significant hits, in which multiple sgRNA against a given human gene were enriched in LCLs that survived NK co-culture, relative to their levels in parallel cultures of LCLs alone. Using a multiple hypothesis test adjusted q-value < 0.05 cutoff, the screen identified 173 genes. CD48 was a top hit, consistent with its known role as an NK-activating receptor highly expressed on LCLs^23,41–43^ (**Fig. 1B**). CD48 is attached to the plasma membrane by a glycosylphosphatidylinositol (GPI) anchor^44,45^, and multiple phosphatidylinositol glycan (PIG) GPI-anchor biosynthesis genes were also top hits (**Fig. 1B-C**), likely due to obligatory roles in CD48 plasma membrane expression. The genes encoding both LFA-1 subunits, ITGAL/CD11A and ITGB2/CD18, as well as multiple factors involved in integrin signaling were also top hits, likely due to roles of LCL/NK adhesion (**Fig. 1B, D**).

Unexpectedly, all subunits of the HUSH complex and its two effectors, which function as an epigenetic silencing system typically of intronless or repetitive DNA elements^46–48^, were identified as top hits (**Fig.1B, D, S1C**). These included the three HUSH recognition module subunits, transcription activation suppressor (TASOR), M-phase phosphoprotein 8 (MPP8, encoded by MPHOSPH8), and periphilin (encoded by PPHLN1) as well as the two HUSH epigenetic effectors, SETDB1 and MORC2. The histone methyltransferase SETDB1 deposits histone 3 lysine 9 trimethylation (H3K9me3) repressive marks at HUSH target sites^48^, whereas the ATP-dependent chromosome remodeler MORC2 restricts chromatin accessibility^49,50^. We will refer to each of these as HUSH complex subunits hereafter.

To validate and extend these results, we repeated the LCL CRISPR screen using CD56+ primary human NK cells, isolated from peripheral blood and activated/expanded by interleukin-15 and NAM treatment^38^. CD48, LFA-1 and HUSH complex subunits were again top screen hits (**Fig. 1E, S1D**). Together, these results indicate that the HUSH complex plays an unexpected role in NK-mediated immune surveillance of EBV-transformed lymphoblastoid B cells.

### LCL HUSH KO confers resistance to NK cell cytotoxicity

We validated that multiple screen hit sgRNAs successfully depleted GM15892 HUSH complex subunits (**Fig. S2A**), further supporting that CRISPR on-target activity was responsible for the screen phenotype. Since CRISPR screen hits can reflect shared pathways, we tested the hypothesis that HUSH might regulate CD48 or LFA1 expression, either directly or indirectly by repressing a repressor. However, plasma membrane CD48 and LFA1-subunit CD11A expression were similar in control versus HUSH subunit KO LCLs (**Fig. S2B**). Similarly, we confirmed that TASOR or MORC2 knockout did not significantly alter LCL proliferation (**Fig. S2C**).

We therefore directly measured effects of LCL HUSH KO, and for comparison CD48 KO, on NK killing. To do so, we used flow cytometry to compare the numbers of Cas9+ GM15892 expressing control, CD48 or HUSH sgRNAs after co-culture for 24 hours with NK at a 2:1 effector to target (E:T) ratio. LCL CD48 or HUSH subunit KO conferred a similar degree of protection from lysis by either YT-INDY or by IL15/NAM-activated primary human NK (**Fig. 2A-B**). TASOR or MORC2 knockout similarly protected a second LCL, GM12878, from attack by YT-INDY, NK-92 or primary human NK cells (**Fig. S3A**).

**Figure 2.**
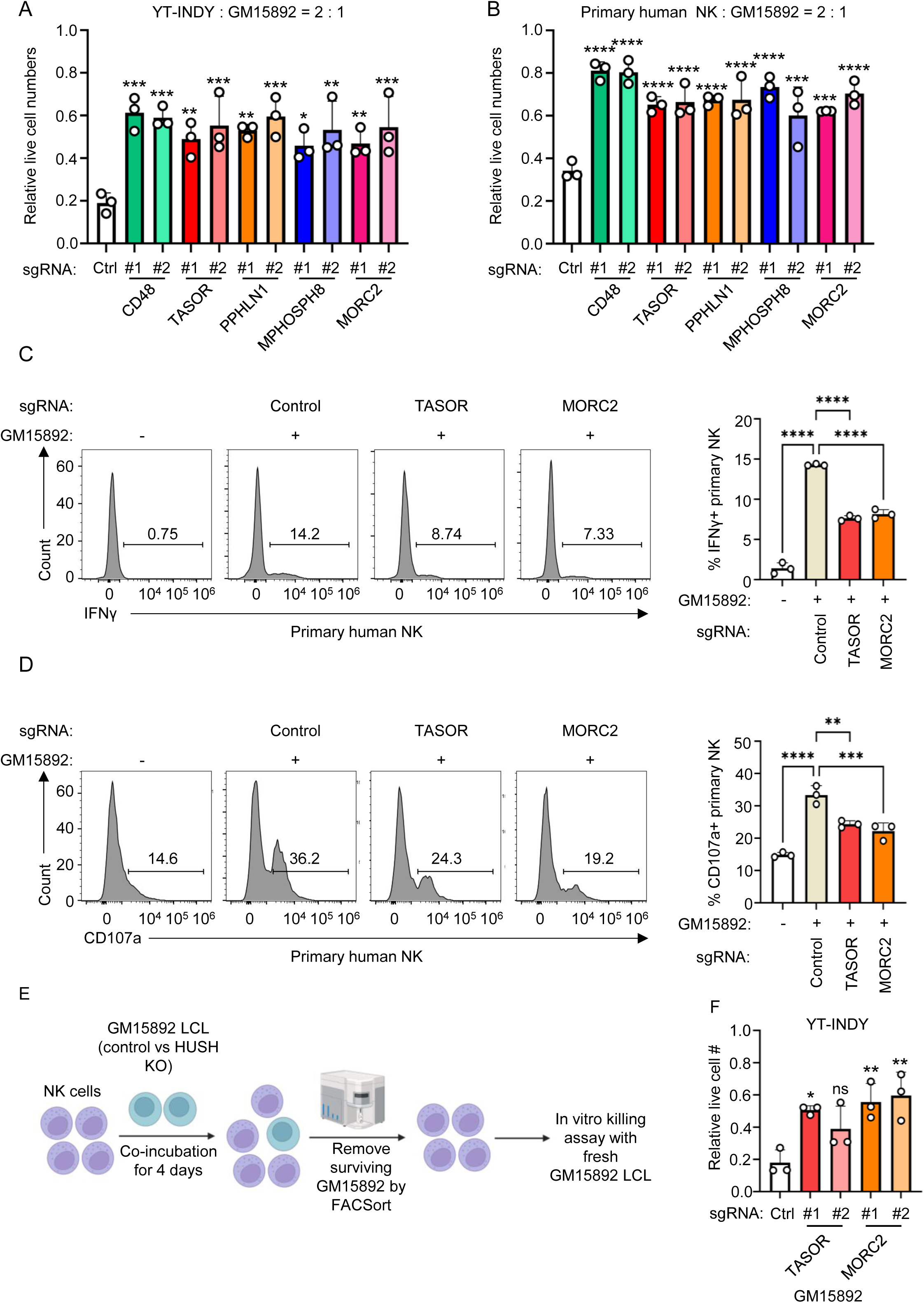
HUSH KO Effects on lymphoblastoid B cell NK surveillance. A) CD48 or HUSH KO effects on LCL killing by YT-INDY. Shown are the mean + SD relative live GM15892 numbers following co-culture with YT-INDY at a 2:1 effector:target (E:T) ratio for 24 hours, from n=3 replicates. Values were normalized by numbers of GM15892 cultured in the absence of YT-INDY. B) CD48 or HUSH KO effects on LCL killing by primary human NK. Shown are the mean + SD relative live GM15892 numbers following co-culture with IL-15/NAM treated primary human NK at a 2:1 E:T ratio for 24 hours, from n=3 replicates. Values were normalized by numbers of GM15892 cultured in the absence of primary NK. C) HUSH KO effects on NK IFNγ^+^ expression. FACS analysis of intracellular IFNγ^+^ expression in primary human IL-15/NAM treated NK cells, cultured alone or co-cultured with Cas9+ GM15892 LCLs expressing the indicated sgRNAs and treated with protein transport inhibitor for 5 hours. Mean + SD values from 3 replicates are shown on the right. D) HUSH KO effects on NK CD107a expression. FACS analysis of plasma membrane CD107a expression in primary human IL-15/NAM treated NK cells, cultured alone or co-cultured with Cas9+ GM15892 LCLs expressing the indicated sgRNAs for 5 hours. Mean + SD values from 3 replicates are shown on the right. E) NK assay schematic. To assess effects of LCL HUSH KO on subsequent NK killing, YT-INDY were co-incubated with control or HUSH KO LCLs for 4 days. NK were FACSorted and then used for killing assays with fresh GM15892 LCL targets. F) Effects of exposure to HUSH KO LCL on subsequent NK killing. NK were pre-incubated with Cas9+ GM15892 with the indicated sgRNA for 4 days, and then used for killing assay with fresh control LCL targets. Mean + SD live cell numbers from n=3 replicates of GM15892 following co-culture with YT-INDY 9 (left) or NK-92 (right) at a 2:1 E:T ratio for 24 hours. Values were normalized by live cell values of GM15892 cultured in the absence of NK. Statistical significance was assessed by one-way ANOVA followed by Tukey’s multiple comparisons test (A-F). *P < 0.05, **P < 0.01, ***P<0.001, ****P < 0.0001, ns, not significant.

We next assessed TASOR KO effects on NK cytotoxicity against two closely matched pairs of EBV+ Burkitt B cells that differ by EBV latency programs. Interestingly, while Jijoye cells with the fully transforming EBV latency III program shared by LCLs were killed by YT-INDY or NK-92 cells, the P3HR-1 subclone, which has an EBV genomic deletion that removed EBNA2 and that converted the viral latency program to a more restricted program^51^, were refractory to NK effects. Similar results were observed with MUTU I vs III cells, which differ only by EBV latency I vs III (EBNA1 is the only EBV protein expressed in latency I)^52^. TASOR knockout significantly protected Jijoye and MUTU III from NK, albeit to a greater degree with YT-INDY than with NK92 (**Fig. S3B**). These results suggest that EBV latency III heightens B cell sensitivity to NK surveillance in a HUSH dependent manner.

To more broadly examine HUSH perturbation effects on EBV-uninfected B-cell sensitivity to NK, we derived a panel of control vs TASOR KO pairs from the diffuse large B-cell lymphoma (DLBCL) lines HBL1, SUDHL5, and SUDHL6, the mantle cell lymphoma lines KARPAS and MAVER-1, and the pre-B acute lymphoblastic leukemia cell line REH. Intriguingly, HUSH KO partially protected HBL1 and MAVER-1 from co-cultured NK cells (**Fig. S3B**). We also tested TASOR KO effects on chronic myelogenous leukemia K562 cells, which lack MHC class I^53,54^ and therefore activate NK. TASOR KO significantly protected K562 from NK-92 effects (K562 were resistant to YT-INDY, as previously identified^55^, **Fig. S3B**). These results indicate that the influence of the HUSH complex over NK surveillance extends beyond EBV-infected cells or B-cells.

### LCL HUSH complex knockout impairs NK cell activation

NK cell-mediated killing involves target cell adhesion, activation, and secretion of cytotoxic granules and/or expression of death receptor ligands^34,56–59^. We therefore next interrogated HUSH perturbation effects on each of these stages. TASOR or MORC2 KO did not significantly inhibit primary NK or YT-INDY adhesion to LCLs (**Fig. S4A-D**). However, co-incubation with GM15892 significantly increased primary human NK intracellular IFN-γ and plasma membrane CD107a levels, hallmarks of NK activation (**Figure 2C-D**). LCL TASOR or MORC2 KO attenuated these NK responses. Similar results were observed when YT-INDY cells were co-cultured with control versus HUSH KO GM15892 or GM12878 LCLs (**Figure S5A-B**), indicating that target LCL HUSH deficiency impairs NK cell activity. By contrast, HUSH perturbation did not render LCLs generally resistant to either apoptosis or necroptosis induction (**Fig. S5C**), suggesting that HUSH KO confers selective resistance to NK-mediated rather than generalized cell death pathways.

To test whether exposure to HUSH KO LCLs impaired subsequent NK responses, we pre-incubated YT-INDY or NK-92 with control, TASOR, or MORC2 KO LCLs for four days. We then sorted NK cells and assessed their cytotoxicity against fresh unedited LCL targets (**Fig. 2E-F**). Pre-exposure to HUSH KO LCL significantly diminished YT-INDY and NK-92 cytotoxicity against LCL targets (**Fig. 2F and S5D**). These results indicate that HUSH KO delivers a negative signal to co-cultured NK cells, which can be sustained even after HUSH KO cell removal.

### HUSH complex represses transcription of PCDHG

The HUSH complex represses target DNA^46–48^, which typically include intronless elements. HUSH represses retroviral and endogenous DNA elements, including LINE-1 and endogenous retroviruses^47,48,60–62^. We therefore hypothesized that HUSH KO inhibited NK surveillance through upregulation of an inhibitory membrane bound or secreted ligand. To test this, we performed Perturb-seq to define TASOR or MORC2 KO effects on GM15892 gene expression. HUSH KO did not uniformly alter EBV gene expression across both TASOR and MORC2 KO cells, though did upregulate multiple endogenous elements (**Fig. S6A-B**). Notably, 30 host B cell genes were up-or down-regulated by both TASOR and MORC2 knockout at an adjusted p-value < 0.05, ≥2-fold change cutoff (**Figure 3A**). Enrichment analysis highlighted that HUSH KO upregulated genes encoding plasma membrane proteins, with many involved in cell adhesion (**Fig. 3B-C**). While none encoded known NK inhibitory ligands, we noted that HUSH KO upregulated multiple protocadherin gamma (PCDHG) genes. Heatmap analysis further highlighted that multiple PCDHG subfamily A and B genes were significantly upregulated by TASOR and/or by MORC2 knockout (**Fig. 3D**). These observations in human cells are reminiscent of previous data, where murine *Mphosph8* or *Morc2* neuronal KO significantly increased expression of multiple protocadherin genes, including of the PCDHG cluster^63,64^.

**Figure 3.**
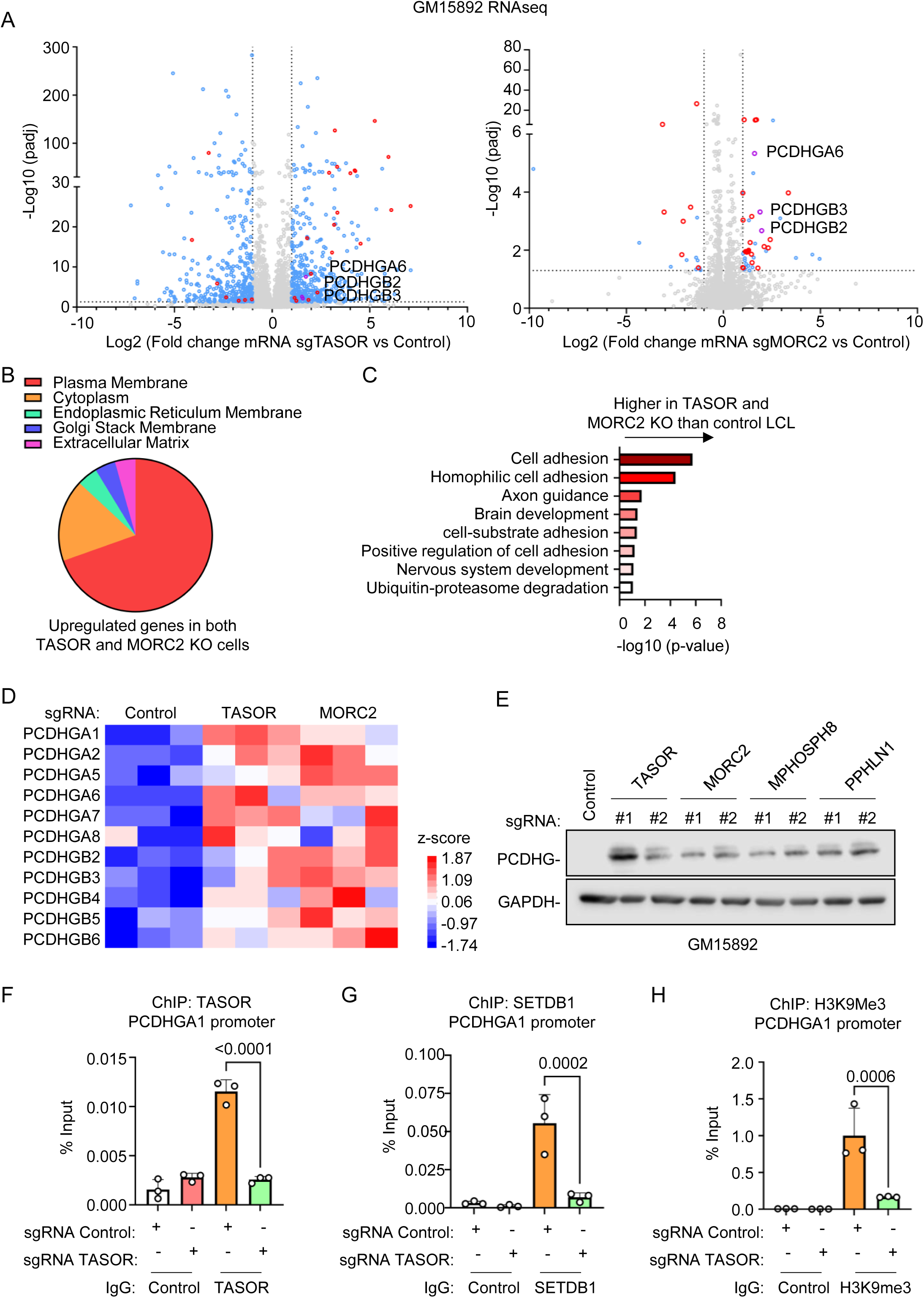
HUSH represses LCL proto-cadherin gamma (PCDHG) expression. A) Analysis of TASOR or MORC2 KO effects on the LCL transcriptome. Shown are volcano plot analyses from n=3 RNAseq replicates of GM15892 that expressed control versus TASOR sgRNAs (left panel) or control versus MORC2 sgRNAs (right panel). Genes with significantly different expressions in both TASOR and MORC2 knockout cells are highlighted in red. PCDHG genes are highlighted in purple. B) GO enrichment analysis of GM15892 LCL genes that were significantly upregulated by both TASOR and MORC2 knockout. C) KEGG analysis of GM15892 LCL genes that were significantly upregulated by both TASOR and MORC2 knockout. D) Heatmap analysis of PCDHG family member mRNA expression in GM15892 LCLs with control, TASOR or MORC2 sgRNAs. Shown are z-score values for family members upregulated by both TASOR and MORC2 knockout. E) Immunoblot analysis of WCL from GM15892 LCLs that expressed the indicated sgRNAs. Blots are representative of n=3 experiments. F-H) ChIP-qPCR analysis of the GM15892 LCL *PCDHGA1* locus. Mean + SD values from n=3 ChIP-qPCR replicates of TASOR (F) or SETDB1 (G) occupancy or of H3K9me3 (H) levels at the *PCDHGA1* promoter region. Anti-TASOR, SETDB1 and H3K9me3 ChIP were performed on chromatin from GM15892 LCLs that expressed the indicated control or TASOR sgRNAs, followed by qPCR. Statistical significance was assessed by two-tailed unpaired Student’s t test (F-H).

Cadherins are a large superfamily of cell surface proteins best known for mediating calcium-dependent cell–cell adhesion integrity^65^, but E-and N-cadherins can serve as NK inhibitory ligands that bind to the inhibitory receptor KLRG1 to suppress NK cell cytotoxicity and cytokine expression^66,67^. In contrast, PCDHGs are structurally distinct and have not been implicated in NK repression. Therefore, to further investigate potential PCDHG NK inhibitory roles, we first validated HUSH KO effects on LCL PCDHG expression on the protein level. Using an antibody against the common PCDHG C-terminal domain, we showed that KO of HUSH subunits encoded by TASOR, MPHOSPH8 or PPHLN1, or KO of the MORC2 effector each derepressed PCDHG protein expression in both GM15892 and GM12878 LCLs (**Figures 3E and S7A**).

To then examine a potential HUSH role in epigenetic repression of LCL PCDHG expression, we first analyzed ENCODE project LCL ChIP-seq datasets^68–70^, which revealed abundant H3K9me3 marks at the GM12878 *PCDHGA* and *PCDHGB* gene clusters, but not at the C3-5 *PCDHGC* exons (**Fig. S7B**). To further characterize potential HUSH roles in LCL PCDHG regulation, we performed ChIP on chromatin purified from GM15892. ChIP-qPCR analysis demonstrated occupancy of TASOR and SETDB1 (the methyltransferase effector of HUSH responsible for H3K9me3 deposition) occupancy, as well as H3K9me3 deposition at the LCL *PCDHGA1* and *PCDHGA2* promoters (**Fig. 3F-H, S7C-E**). TASOR knockout significantly reduced PCDHGA1 and PCDHGA2 promoter H3K9me3 marks (**Fig. 3F-H, S7C-E**). These findings suggest that HUSH epigenetically represses LCL PCDHG expression.

### PCDHG is necessary for HUSH KO Protection from NK

We next tested the hypothesis that PCDHG expression was involved in the inhibitory effects of LCL HUSH KO on NK. For these experiments, we knocked out PCDHG in LCLs by targeting an exon that encodes a C-terminal tail region shared by all PCDHG family members^63^. We then introduced control, TASOR or MORC2 targeting sgRNAs in GM15892. Immunoblot analysis confirmed PCDHG expression in TASOR or MORC2 KO LCLs, but not in those with double PCDHG KO (**Figure 4A**). Importantly, PCDHG knockout by either of two independent sgRNAs significantly increased GM15892 LCL killing by YT-INDY, to levels similar to those observed in control LCLs (**Figure 4B**). Similar results were observed in GM12878 challenged by YT-INDY or NK-92 (**Fig. S8A-C**).

**Figure 4.**
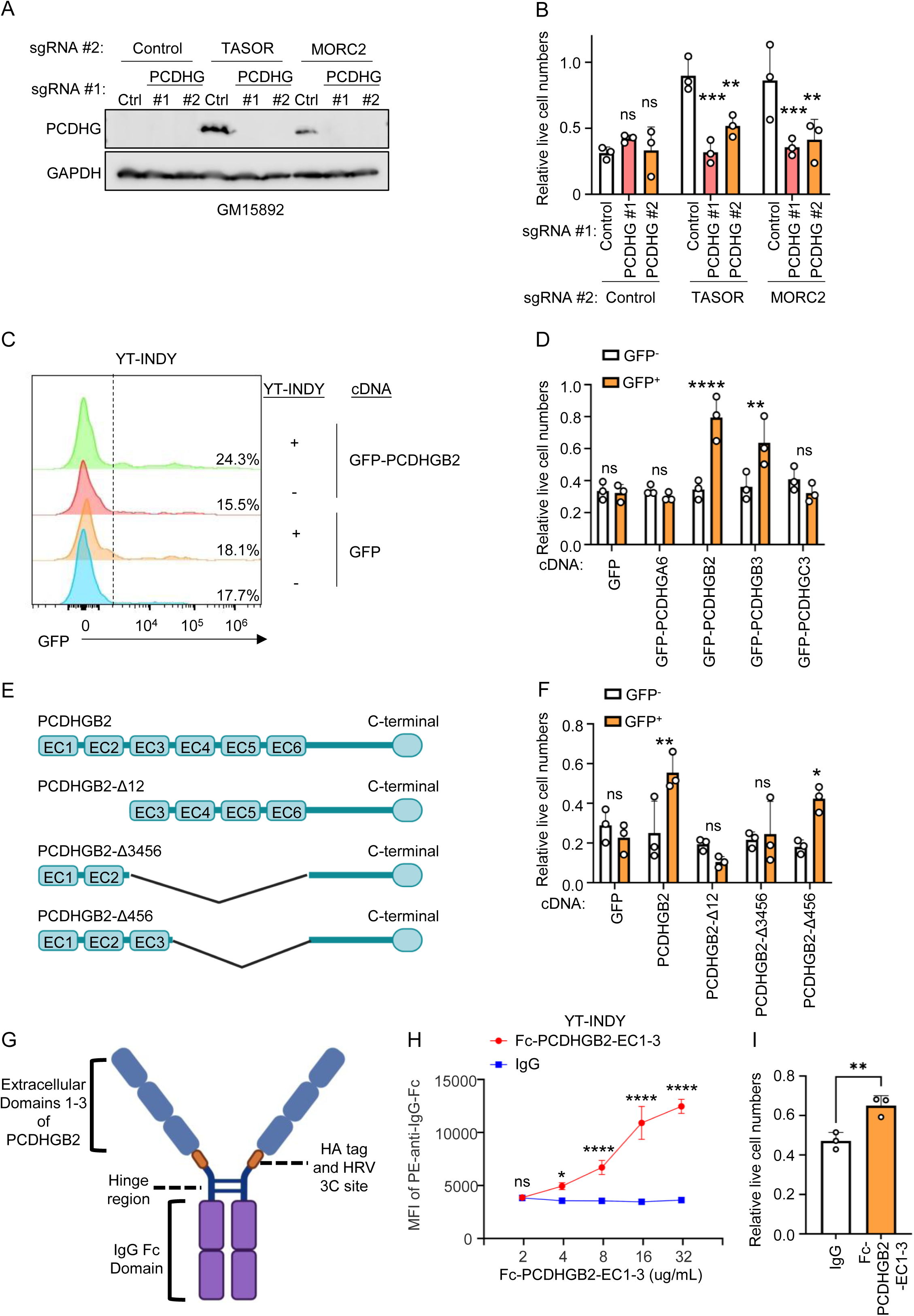
LCL PCDHG expression inhibits NK lysis. A) Immunoblot analysis of WCL from GM15892 that expressed the indicated sgRNAs targeting control, TASOR or MORC2, together with sgRNAs targeting an exon that encodes a common PCDHG C-terminal domain. A pan-PCDHG antibody was used for the blot. B) Analysis of PCDGH roles in HUSH KO LCL protection from NK lysis. Shown are the mean + SD values from n=3 replicates of GM15892 LCLs NK lysis assays. Cas9+ GM15892 expressed the indicated control or HUSH sgRNAs, together with control sgRNA or with an sgRNA targeting a common *PCDHG* exon. LCLs were co-cultured with YT-INDY at 2:1 E:T ratio for 24 hours. Live cell LCL values were normalized by those of identical GM15892 cultured in the absence of YT-INDY. C) LCL PCDHGB2 expression effects on protection from NK lysis. Representative FACS plot from n=3 replicates of GFP expression in GM15892 LCLs electroporated with either empty vector or vector encoding GFP-tagged PCDHG, following co-culture with YT-INDY cells at an effector-to-target (E:T) ratio of 2:1 for 24 hours, or cultured alone (control). GM15892 were stained with CellTrace Violet to distinguish them from YT-INDY cells. D) Effects of PCDHG family member expression on LCL protection from NK lysis. Mean + SD relative live cell values from n=3 replicates of GM15892 LCLs that were electroporated with the indicated DNA and then co-cultured with YT-INDY for 24 hours at an E:T of 2:1. Live cell values were normalized by those of identical LCLs cultured in the absence of NK. E) Schematic diagram illustrating PCDHGB2 domains in wildtype versus PCDHGB2 truncation mutants. F) Effects of wildtype versus PCDGHB2 deletion mutant expression on LCL protection from NK lysis. Mean + SD relative live cell values from n=3 replicates of GM15892 LCLs that were electroporated with the indicated PCDHB2 constructs and then co-cultured with YT-INDY for 24 hours an E:T of 2:1. Live cell values were normalized by those of identical LCLs cultured in the absence of NK. GM15892 cells were stained with CellTrace Violet to distinguish them from YT-INDY. G) Schematic diagram illustrating the recombinant Fc-PCDHGB2-EC1-3 fusion protein. H) FACS analysis of YT-INDY labeling by the Fc-PCDHGB2-EC1-3 fusion protein. Shown are mean ± SD mean fluorescence intensity (MFI) values of from n=3 replicates of YT-INDY co-incubated with the indicated concentrations of control IgG versus Fc-PCDHGB2-EC1-3 for 3 hours at 37 degrees prior to labeling with secondary PE-tagged anti-IgG and FACS analysis. I) Fc-PCDHGB2-EC1-3 effects on NK-mediated LCL lysis. Shown are mean + SD relative live cell values of GM15892 cells co-incubated with YT-INDY cells that were preincubated with 16 ug/mL Fc-PCDHGB2-EC1-3 protein or with control IgG for 24 hours. YT-INDY were then co-incubated with LCLs at a 2:1 E:T ratio for 24 hours. LCL live cell values from cultures with Fc-PCDHGB2-EC1-3 were normalized by values from cultures of IgG treated NK. Statistical significance was assessed by two-tailed unpaired Student’s t test (D,F,I) or two-way ANOVA followed by Tukey’s multiple comparisons test (B,H). *P < 0.05, **P < 0.01, ***P<0.001, ****P < 0.0001, ns, not significant.

We next tested whether expression of individual PCDHGs were sufficient to protect LCL from NK attack. Expression of either PCDHGB2 or PCDHGB3, whose levels are significantly increased by HUSH KO, significantly increased the numbers of live GM15892 LCLs after co-culture with YT-INDY for 24 hours (**Fig. 4C-D, S8D-E**). In contrast, PCDHGA6 or PCDHGC3 overexpression did not confer this resistance, potentially suggesting that specific PCDHGs exert inhibitory effects, or perhaps that these PCDHGs were not expressed at the same levels at the plasma membrane in our transient expression system (**Fig. 4C-D**).

The PCDHG extracellular region is comprised of six N-terminal extracellular cadherin (EC) domains. To determine which EC domains were critical for NK resistance, we generated a series of PCDHGB2 truncation mutants, deleted for different combinations of EC domains (EC1-1, EC3-6 or EC4-6, **Fig. 4E**). We confirmed that each was expressed and that each trafficked to the LCL plasma membrane (**Fig. S8F**). Overexpression of the PCDHGB2 ΔEC4-6 mutant conferred GM15892 protection from YT-INDY killing, whereas overexpression of ΔEC1-2 or ΔEC 3-6 PCDHGB2 mutants failed to do so. These results indicate that the PCDHG EC1-3 domains are required for protection from NK lysis (**Fig. 4F**).

Based on these findings, we constructed, expressed and purified a recombinant chimera of the PCDHGB2 EC1–3 domains with a C-terminal HA-tag and IgG Fc (fragment crystallizable) domain (**Fig. 4G and S8G**). Flow-cytometry-based immunostaining experiments revealed that the PCDHGB2-EC1–3-Fc fusion protein could specifically label YT-INDY and NK-92, but not HEK293T, in a dose-dependent manner (**Fig. 4H and S8H-I**). Moreover, preincubation of YT-INDY or NK92 cells with the PCDHG-EC1-3-Fc-fusion protein for 24 hours significantly reduced their cytotoxicity toward GM15892 LCLs (**Fig. 4I and S8J**). Collectively, these results suggest that PCDHG EC1-3 interacts with a specific NK cell plasma membrane inhibitory receptor to impair NK surveillance.

### PCDHG inhibits NK cell activity through NKG2A/KLRC1 engagement

To identify the PCDHG NK counter receptor, we performed a CRISPR screen with a sgRNA library which targets most human transmembrane protein encoding genes with 10 independent sgRNAs^71,72^. Cas9+ YT-INDY cells were transduced at a multiplicity of infection of 0.3, puromycin selected and stained with PE-labelled Fc-PCDHGB2-EC1-3 (16 μg/ml for 3 hours). The 5% of YT-INDY cells with the lowest PE signal (with the least binding to PE-Fc-PCDHB2) were isolated by flow cytometry. PCR-amplified sgRNA sequences from the input library versus from sorted cells were quantitated by next generation DNA sequencing (**Fig. 5A**).

**Figure 5.**
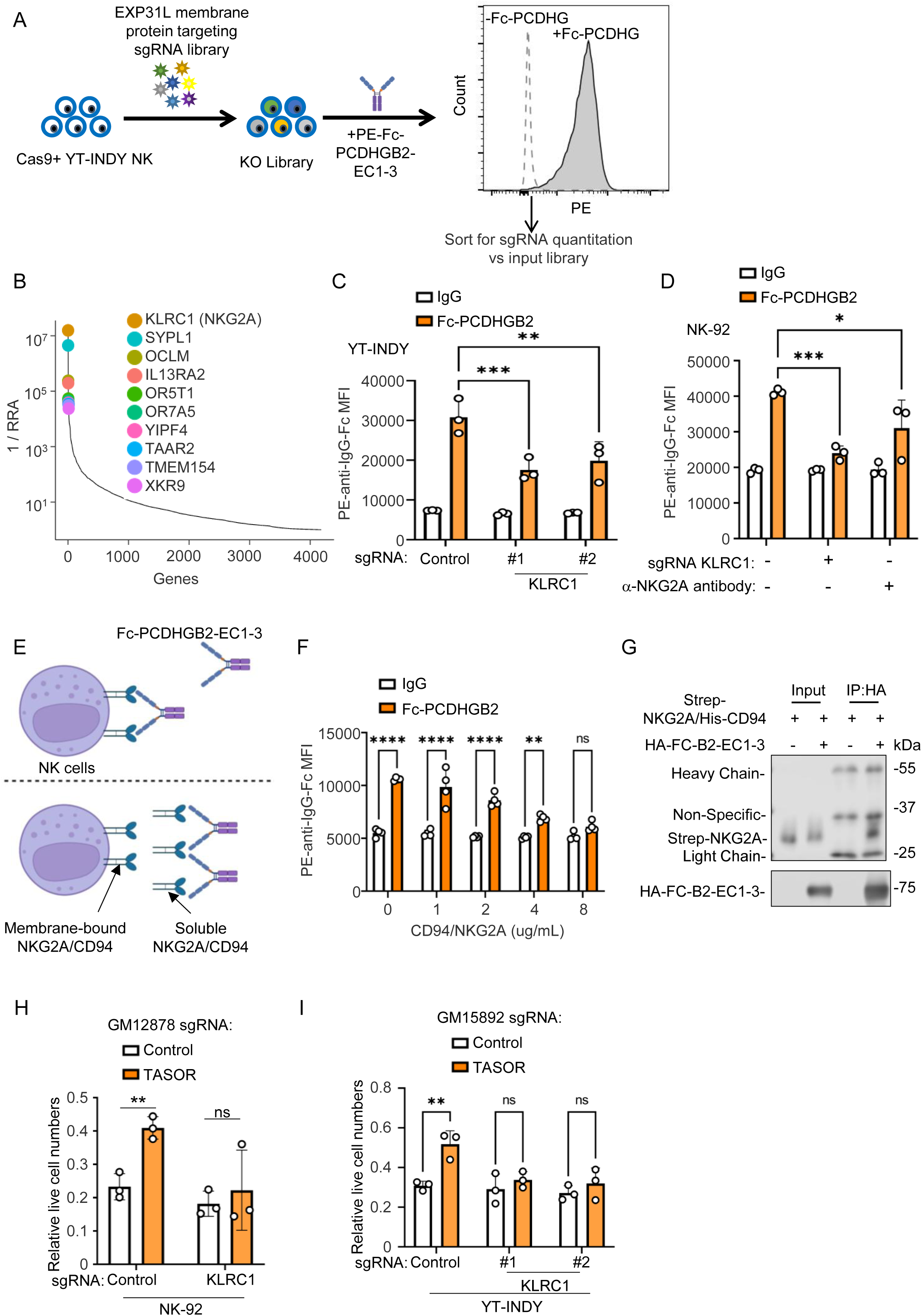
Human genome-wide CRISPR-Cas9 screen for the NK PCDHG counter-receptor. A) NK CRISPR screen workflow. Cas9+ YT-INDY cells were transduced with the membrane protein targeting sgRNA library. The knockout library was stained with PE-tagged Fc-PCDHGB2-EC1-3 (16 μg/mL) for 3 hours. The 5% of cells with the lowest PE-Fc-PCDHGB2-EC1-3 were sorted. sgRNA abundances in unsorted library input vs sorted cells were quantified to enable hit identification. B) Screen hits. MAGeCK analysis for CRISPR screen hits, ranked based on robust rank aggregation (RRA) scores (y-axis) versus genes (x-axis). KLRC1/NKG2A was the top hit. C) KLRC1/NKG2A KO effects on YT-INDY labeling by Fc-PCDHGB2. Cas9+ YT-INDY that expressed control or screen hit KLRC1 sgRNAs were incubated with Fc-PCDHGB2-EC1-3 (16 μg/mL) for 3 hours and then by PE-tagged anti-IgG Fc antibody. Shown are mean + SD MFI values from n=3 replicates. D) Anti-NKG2A antibody effects on NK-92 labeling by Fc-PCDHGB2. Shown are mean + SD MFI values from n=3 replicates of Cas9+ NK92 cells that expressed control or KLRC1 targeting sgRNA or that were pre-treated with the anti-NKG2A antibody Z-199 at 10 μg/mL for 1 hour and that were then incubated with Fc-PCDHGB2-EC1-3 (16 μg/mL) for 3 hours and then by PE-tagged anti-IgG Fc antibody. E) Schematic diagram of the soluble NKG2A/CD94 competition assay for NK labeling by Fc-PCDHGB2-EC1-3. YT-INDY cells were incubated with Fc-PCDHGB2-EC1-3 in the presence or absence of increasing concentrations of soluble NKG2A/CD94. F) NKG2A/CD94 competition for NK labeling by Fc-PCDHGB2-EC1-3. Mean + SD MFI values from n=3 replicates of YT-INDY cells that were incubated with Fc-PCDHGB2-EC1-3 (16μg/mL) together with the indicated concentrations of soluble NKG2A/CD94, as in (E). YT-INDY plasma membrane Fc fusion protein labeling was detected by PE-anti-IgG Fc antibody. G) Co-immunoprecipitation analysis of Fc-PCDHGB2-EC1-3 association with recombinant NKG2A/CD94. Fc-PCDHGB2-EC1-3 and NKG2A/CD94 (5μg) were co-incubated for 4 hours at 4 degrees, and anti-HA magnetic beads were used to purify HA-Fc-PCDHGB2-EC1-3. Shown are immunoblot analysis of 1% input vs anti-HA-tagged Fc-PCDHGB2-EC1-3 immunoprecipitated complexes. H) Analysis of KRLC1/NKG2A depletion effects on NK-92-mediated killing of control vs HUSH KO LCLs. Mean + SD relative live cell values from n=3 replicates of Cas9+ GM12878 LCLs that expressed control or TASOR sgRNAs, and that were co-incubated with Cas9+ NK-92 cells that expressed control or KLRC1 sgRNAs. NK92 and LCLs were co-cultured at a 2:1 E:T ratio for 24 hours. I) Analysis of KRLC1/NKG2A depletion effects on YT-INDY NK-mediated killing of control vs HUSH KO LCLs. Mean + SD relative live cell values from n=3 replicates of Cas9+ GM15892 LCLs that expressed control or TASOR sgRNAs, and that were co-incubated with Cas9+ YT-INDY cells that expressed control or KLRC1 sgRNAs. YT-INDY and LCLs were co-cultured at a 2:1 E:T ratio for 24 hours. Statistical significance was assessed by two-way ANOVA followed by Tukey’s multiple comparisons test (C,D,F,H,I). *P < 0.05, **P < 0.01, ***P<0.001, ****P < 0.0001, ns, not significant.

Using three screen replicates, we used MAGeCK analysis to analyze enriched genes based on robust rank aggregation (RRA) scores. The top hit, KLRC1, encodes the well-characterized NK cell inhibitory receptor NKG2A (**Fig. 5B**). NKG2A typically forms an inhibitory heterodimer together with KLRD1-encoded CD94, which interacts with HLA-E to suppress NK cell activity^73,74^. However, sgRNAs against KLRD1 was not present in the library. NK NKG2A/KLRC1 KO by either of two independent library sgRNAs significantly reduced YT-INDY plasma membrane labeling by PE-PCDHGB2-EC1-3-Fc (**Fig. 5C and S9A**). Likewise, NK92 NKG2A/KLRC1 KO blocked PCDHGB2-EC1-3-Fc plasma membrane labeling, and an anti-NKG2A antibody competed with Fc-PCDHGB2 for NK92 plasma membrane labeling (**Fig. 5D and S9B**).

To further characterize the interaction between PCDHGB2 and NKG2A/CD94, we generated soluble strep-tagged NKG2A and His-tagged CD94 proteins (**Fig. S9C**). The soluble NKG2A/CD94 complex competed with Fc-PCDHGB2-EC1-3 for YT-INDY plasma membrane labeling, in a dose-dependent manner (**Fig. 5E-F**). Further suggesting direct interaction, HA-tagged Fc-PCDHGB2-EC1-3 co-immunoprecipitated soluble NKG2A/CD94 (**Fig. 5G**).

Furthermore, NKG2A expression was necessary for NK92 or YT-INDY inhibition by TASOR KO in GM12878 or GM15892 LCLs (**Fig. 5H-I**). Collectively, these results suggest that PCDHG inhibits NK cell activity through direct interaction with NKG2A/CD94.

### TASOR KO promotes LCL xenograft resistance to NK killing *in vivo*

To assess the impact of HUSH complex perturbation on NK sensitivity *in vivo*, we used NOD.*scid*.*Il2Rγc^null^* (NSG)-IL15 mice, which lack endogenous T and NK cells, but transgenically express human IL-15 to support adoptively transferred human NK cells^75^. Control or TASOR KO GM12878 cells were injected subcutaneously into bilateral flanks to establish LCL xenograft tumors. Once tumors became palpable, 10 million NK-92 cells or PBS were infused intravenously, and tumor growth was monitored (**Fig. 6A**). Adoptively transferred NK cells markedly inhibited the growth of control LCL xenografts, but had significantly less inhibitory effect on TASOR KO tumors (**Fig. 6B-C**). At the endpoint, tumors were harvested, dissociated into single-cell suspension and analyzed by flow cytometry. TASOR KO tumors contained fewer infiltrating CD56⁺ NK cells than control tumors (**Fig. 6D**). Moreover, tumor-infiltrating NK cells isolated from TASOR KO tumors expressed lower IFNγ levels following PMA/ionomycin stimulation than those isolated from control LCL xenograft tumors (**Fig. 6E**).

**Figure 6.**
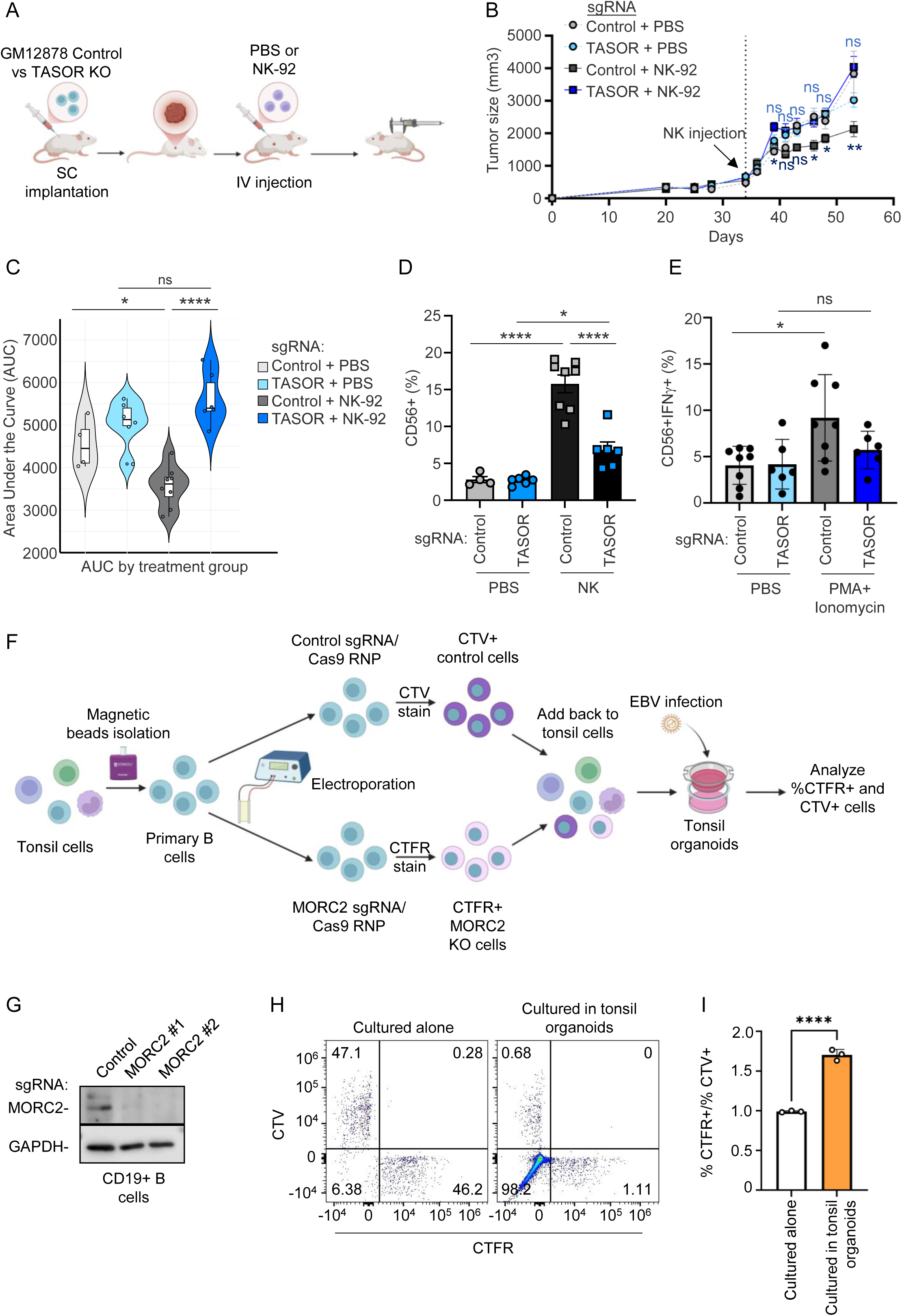
Analysis of TASOR knockout effects on NK surveillance of LCL murine xenografts *in vivo* and in human tonsil organoids. A) Analysis of HUSH effects on LCL NK surveillance *in vivo*. NSG-IL15 mice were subcutaneously injected with 10 million Cas9+ GM12878 LCLs expressing control or TASOR sgRNAs with Matrigel into bilateral mouse flanks (two xenografts/mouse). When xenograft tumors reached 500 mm^3^, 10 million NK-92 cells or PBS vehicle were adoptively transferred intravenously. Effects on tumor sizes were serially measured. Bilateral flank tumors were analyzed from n=3 mice with control tumors/PBS, 4 mice with TASOR KO/PBS, 5 mice with control tumor/NK-92, 5 mice with TASOR KO LCLs/NK-92. B) Mean ± SEM xenograft tumor volumes from mice treated as described in panel (A). NK cells were adoptively transferred on day 35. The asterisks or ns above the curve indicate statistical differences for PBS-injected mice, and those below the curve indicate statistical differences for NK92-injected mice. C) Violin plot analysis of tumor volume area under the curve from the experiment shown in (B), calculated by the RStudio Pracma package. D) Analysis of tumor xenograft NK infiltration. Mean ± SD CD56⁺ NK percentages out of total tumor xenograft cells at the time of tumor explant. E) Analysis of tumor xenograft NK infiltration. Mean ± SD percentages of IFNγ+ NK cells out of total tumor xenograft cells at the time of tumor explant, following PMA (50ng/mL) and ionomycin (1μg/mL) treatment for 4 hours. F) Analysis of MORC2 KO effects on EBV+ B cell NK surveillance within human tonsil organoids. CD19+ B cells isolated by magnetic bead negative selection from human tonsil cell suspensions were electroporated with Cas9 RNP loaded with non-targeting control or MORC2 sgRNA. Control cells were stained with 2mM Celltrace Violet (CTV), MORC2 KO cells were stained with 0.5mM Celltrace Far Red (CTFR). 0.2 million of CTV and CTFR stained cells were mixed at a 1:1 ratio and cultured alone or together with 6 million autologous tonsil cells in transwell plate organoid cultures. Cultures were infected with GFP+ B95.8 EBV at MOI of 0.1. At day 5 post infection, the percentages of EBV+/GFP+ CTV+ and CTFR+ cells were analyzed by FACS. G) Immunoblot analysis of WCL from CD19+ B cells electroporated with the indicated Cas9 RNPs described in (F). Blots are representative of n=3 experiments. H) FACS analysis of CTV-stained control and CTFR-stained MORC2 KO primary B cells cultured in human tonsil organoids as described in F. Plots are representative of n=3 independent replicates. I) Analysis of MORC2 KO effects on EBV+ B cell survival in human tonsil organoids. Shown are mean + SD values of EBV+/GFP+ CTFR+ MORC2 KO versus CTV+ control B cells, from n=3 replicates as in (H). Statistical significance was assessed by one-way ANOVA followed by Tukey’s multiple comparisons test (C-E), two-way ANOVA mixed effects analysis (B) or two-tailed unpaired Student’s t test (I). *P < 0.05, ****P < 0.0001, ns, not significant.

We next assessed the effect of HUSH perturbation on EBV-infected B cells in a recently developed human tonsil organoid model^76^, using EBV seronegative donors. CD19+ B cells were isolated from the tonsil cell admixture and electroporated with Cas9 ribonucleoprotein (RNP) complexes loaded with control or MORC2 targeting sgRNA. Control and MORC2 edited cells were stained with Celltrace Violet (CTV) or Celltrace far red (CTFR), respectively. Labelled cells were mixed and added back to the tonsil cell admixture to establish tonsil organoids (**Fig. 6F**). B cell CRISPR editing was confirmed by immunoblot (**Fig. 6G**). Tonsil organoids were infected with the EBV B95.8 strain at a 0.1 multiplicity of infection. Five days later, the percentages of CTV+ control versus MORC2 KO CTVR+ cells were quantitated by FACS. As expected, we observed equal numbers of EBV-infected CTFR+ MORC2 KO versus CTV+ control cells when cultured alone. By contrast, the ratio of CTFR+ MORC2 KO to CTV+ control B cell ratio was substantially increased after five days of culture within tonsil organoids (**Fig. 6H-I),** suggesting that there was a selection against CTV+ B-cells in this microenvironment. These results support the hypothesis that HUSH KO enhances the resistance of EBV-infected B-cells to NK attack in a human secondary lymphoid organ-like microenvironment.

### PCDHG protects follicular lymphoma and neuroblastoma cells from NK attack

We wondered whether PCDHG expression might also be used by EBV-negative tumors to promote NK resistance. Notably, several small studies have found elevated PCDHG expression in follicular lymphoma (FL) tumors^77,78^, a common EBV-negative non-Hodgkin lymphoma. In one such study, 19 of 20 FL grade 1-2 samples and 5 of 9 grade 3A FL samples over-expressed PCDHG^4^. To gain further insights, we tested the functional significance of PCDHG expression in a human tumor derived FL cell line, RL^79^. Confocal microscopy analysis confirmed that RL express PCDHG (**Fig. 7A**). In parallel, immunoblot analysis revealed that RL cells express lower levels of multiple HUSH subunits compared to GM12878 LCLs, consistent with the possibility that reduced HUSH activity may underlie the elevated PCDHG expression observed in RL cells (**Fig. 7B**). We next tested whether PCDHG expression protected RL from NK cell-mediated cytotoxicity. We derived control versus PCDHG knockout RL cells by targeting a common PCDHG exon shared by all family members (**Fig. 7C**). We co-cultured control versus PCDHG KO RL with NK-92 or with IL-15/NAM activated primary human NK cells for 24 hours at a 2:1 effector:target ratio. PCDHG depletion significantly increased RL killing both by types of NK cells (**Fig. 7D-E**). Similarly, PCDHG KO by one of two sgRNAs targeting common exons increased degranulation of NK-92 that were co-cultured with RL cells for 24 hours, indicative of increased NK activation (**Fig. 7F**).

**Figure 7.**
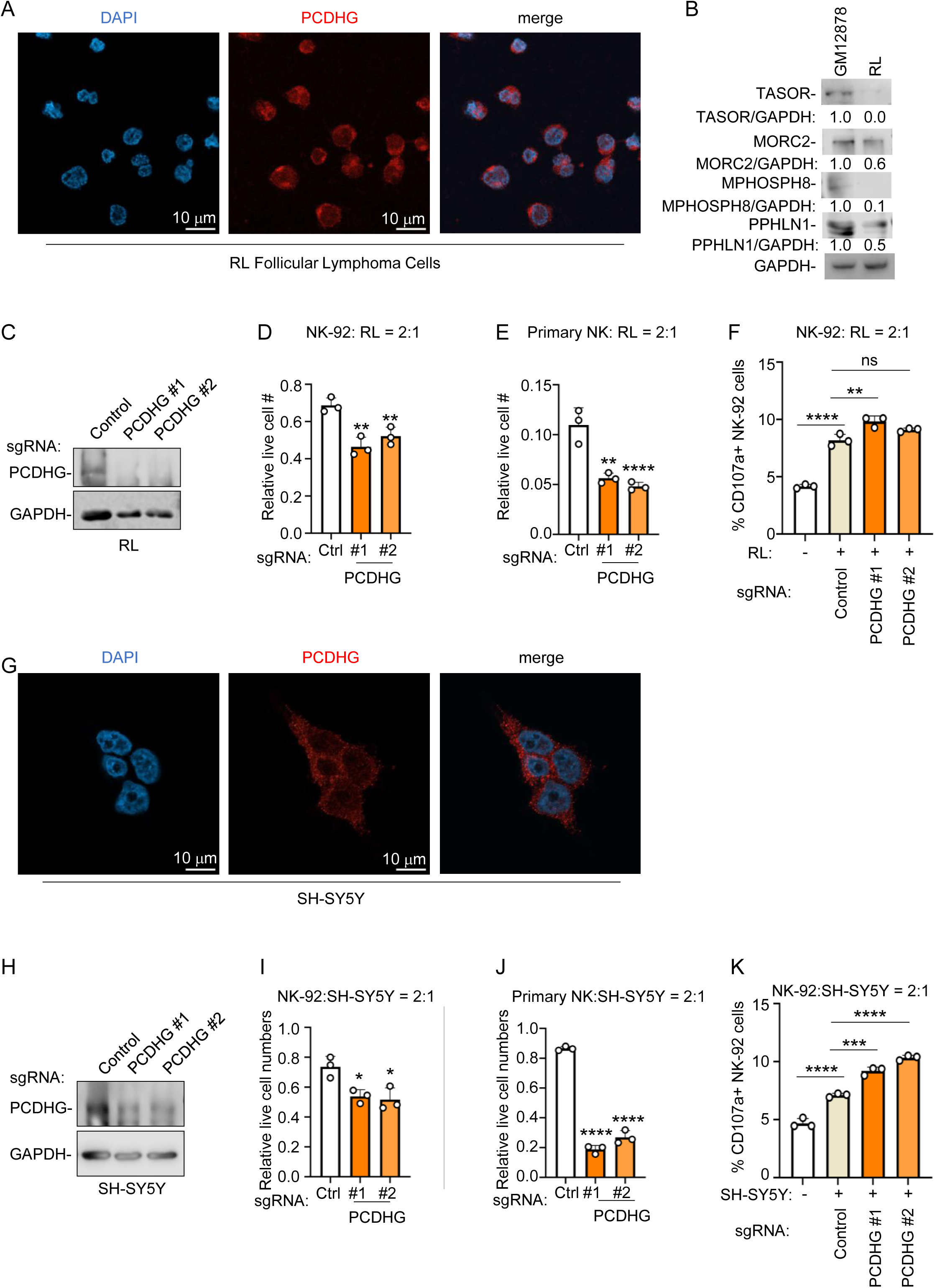
**Analysis of PCDHG knockout effects on NK responses to follicular lymphoma and neuroblastoma tumor cells.**A) Analysis of RL follicular lymphoma PCDHG expression. Representative confocal immunofluorescence images of RL cell PCDHG expression. B) Immunoblot analysis of WCL from GM12878 LCLs versus from RL follicular lymphoma cells. Relative GAPDH-normalized densitometry ratios are shown, with LCL levels set to 1. C) Immunoblot analysis of WCL from Cas9+ RL cells that expressed control or an sgRNA targeting a common *PCDHG* exon. D) Analysis of PCDHG KO on NK-92 mediated RL killing. Mean + SD relative live cell values from n=3 replicates of Cas9+ RL cells that expressed control or PCDHG sgRNAs, and that were co-cultured alone or with NK-92 at a 2:1 E:T ratio for 18 hours. Shown are the ratios of live cells from RL cultured with NK vs alone. E) Analysis of PCDHG KO on primary human NK mediated RL killing. Mean + SD relative live cell values from n=3 replicates of Cas9+ RL cells that expressed control or PCDHG sgRNAs, and that were co-cultured alone or with IL-15/NAM treated primary human NK at a 2:1 E:T ratio for 18 hours. Shown are the ratios of live cells from RL cultured with NK vs alone. F) Analysis of RL PCDHG KO on NK-92 CD107a levels. Mean + SD percentages of CD107a+ NK-92 following co-cultured for 5 hours with Cas9+ RL cells that expressed control versus PCDHG sgRNAs. G) Analysis of SH-SY5Y neuroblastoma PCDHG expression. Representative confocal immunofluorescence images of SH-SY5Y PCDHG expression. H) Immunoblot analysis of WCL from Cas9+ SH-SY5Y cells that expressed control or an sgRNA targeting a common *PCDHG* exon. I) Analysis of PCDHG KO on NK-92 mediated SH-SY5Y killing. Mean + SD relative live cell values from n=3 replicates of Cas9+ SH-SY5Y cells that expressed control or PCDHG sgRNAs, and that were co-cultured alone or with NK-92 at a 2:1 E:T ratio for 18 hours. Shown are the ratios of live cells from SH-SY5Y cultured with NK vs alone. J) Analysis of PCDHG KO on primary human NK mediated SH-SY5Y killing. Mean + SD relative live cell values from n=3 replicates of Cas9+ SH-SY5Y cells that expressed control or PCDHG sgRNAs, and that were co-cultured alone or with IL-15/NAM treated primary human NK at a 2:1 E:T ratio for 18 hours. Shown are the ratios of live cells from SH-SY5Y cultured with NK vs alone. K) Analysis of SH-SY5Y PCDHG KO on NK-92 CD107a levels. Mean + SD percentages of CD107a+ NK-92 following co-cultured for 5 hours with Cas9+ SH-SY5Y cells that expressed control versus PCDHG sgRNAs. Statistical significance was assessed by one-way ANOVA followed by Tukey’s multiple comparisons test (D-F,I-K). *P < 0.05, **P < 0.01, ***P<0.001, ****P < 0.0001, ns, not significant.

Finally, since neurons highly express PCDHG, we hypothesized that neuronal PCDHG expression has been co-opted as a “don’t kill me” signal to protect particular neuroblastoma tumors. To test this hypothesis, we utilized the human neuroblastoma cell line SH-SY5Y, which we found to abundantly express PCDHG (**Fig. 7G**). As above, we used CRISPR to establish control versus PCDHG depleted SH-SY5Y by targeting a common *PCDHG* exon with either of two sgRNAs (**Fig. 7H**). PCDHG depletion by either of the two sgRNA sensitized SH-SY5Y to killing by NK-92 or IL-15/NAM activated primary human NK cells (**Fig. 7I-J**), and increased NK-92 CD107a plasma membrane levels (**Fig. 7K**). These results demonstrate that both follicular lymphoma and neuroblastoma can exploit PCDHG expression to evade NK-mediated immune surveillance, extending the relevance of the HUSH-PCDHG-NKG2A axis beyond EBV-driven lymphoproliferative disease.

## Discussion

NK cells can recognize and kill EBV-immortalized B-cells, raising the question of how virally transformed cells such as PTLD escape NK surveillance to form tumors. Here, we tested the hypotheses that EBV-transformed lymphoblastoid B cells can evade NK detection through inactivation of specific host genomic loci. Human genome-wide CRISPR/Cas9 screens identified that the HUSH complex is an important mediator of LCL sensitivity to NK immune recognition.

Mechanistically, transcriptomic analyses showed that HUSH inactivation de-repressed PCDHG expression, and functionally PCDHG expression conferred protection from NK lysis both *in vitro* and *in vivo*. An orthogonal CRISPR screen identified NKG2A as a candidate NK counter-receptor for PCDHG. This latter finding was confirmed by 1) direct binding studies using purified receptor and ligand proteins; 2) CD94/NKG2A-specific mAb blocking experiments. Collectively, these studies identify a pathway by which EBV-transformed lymphoblastoid cells and other cancers can inhibit NK surveillance through cognate PCDHG/NKG2A/CD94 interactions (**Fig. S10**).

Our results suggest that the HUSH complex plays two roles in innate immunity. First, it represses foreign DNA elements^80–82^, including retroviruses and lentiviruses^83–89^, herpes simplex virus^90^, hepatitis B virus^91^, adeno-associated virus^92,93^ and endogenous retroviral, mobile and repetitive DNA elements^47,80,94–96^. Second, it regulates cellular sensitivity to NK surveillance by modulating the expression of PCDHG genes that are ligands for the CD94/NKG2A inhibitory receptor.

The known ligand for the CD94/NKG2A heterodimer is the non-classical Class I molecule HLA-E, which binds peptides derived from the leader sequence of classical Class I MHC molecules^97–99^. To our knowledge, the demonstration that CD94/NKG2A can also bind γ-proto-cadherins to protect target cells from NK cell lysis is a rare example of an inhibitory receptor able to bind two structurally unrelated ligands. This finding contrasts with data showing that many NK activating receptors can bind several structurally unrelated ligands^100^. For example, the NKp30 receptor (NCR3, CD337) is able to bind the human Ig-SF protein B7-H6^101^, poxvirus haemagglutinins^102^, the Duffy binding-like 1-α domain of Plasmodium falciparum erythrocyte membrane protein-1^103^ and β-1,3 glucans in the fungal cell walls of *Cryptococcus neoformans* and *Candida albicans*^104^. This degeneracy of ligand recognition is presumably not harmful, because in all these contexts, the recognition of these various ligand signals danger. The evolutionary advantage of having multiple ligands for a single inhibitory receptor is not known, but may be related to the observation that MHC class I proteins are only expressed by specific subsets of CNS neurons and the expression level of these molecules is strongly regulated by neural activity^105,106^. An intriguing possibility is that CD94/NKG2A recognition of PCDHG evolved as a means to avoid NK attack of sensitive neuronal tissues. Whereas NK cells are typically excluded from the central nervous system, they can cross the blood-brain barrier in neuropathological conditions^107–109^. PCDHG are expressed in most neurons, and play obligatory roles in neuronal survival^110,111^. PCDHG are found at synaptic clefts and are enriched within postsynaptic density fractions of synaptosomes^110–113^. PCDHG play key roles in control of neuronal synapse formation and touch sensory neuron peripheral branching and synaptic development^63^. Nonetheless, HUSH inactivation within the nervous system further de-represses proto-cadherin expression and alters brain architecture, impairs motor function and reduces lifespan^64^. Twenty one of 55 genes upregulated by MPHOSPH8 inactivation were proto-cadherins^64^. NK may have evolved to recognize and leverage this defining neuronal feature through NKG2A/CD94 as a ‘don’t kill me’ signal, setting a high threshold for NK activation within the CNS. At the same time, however, we now propose that this defense mechanism can be subverted by tumors and virally infected cells so that HUSH disruption leads to protection from NK cell lysis.

We note that previous work reported a YT-INDY variant lacking CD94/NKG2A^114^. However, our YT-INDY stock consistently expresses CD94/NKG2A by flow cytometry. NK-like cell lines are known to undergo phenotypic drift during long-term laboratory propagation, and differences in receptor expression across laboratories have been documented for multiple NK-derived lines. Thus, the discrepancy likely reflects subclone divergence rather than technical variation.

How might EBV infected tumors, and more generally other tumors including FL and neuroblastoma subvert HUSH to de-repress PCDHG? Tumor genome sequencing has not yielded evidence of frequent inactivating mutations of HUSH, so we hypothesize that FL may epigenetically alter the levels of HUSH subunits such as SETDB1 or MORC2 to reach a threshold at which PCDHG expression becomes de-repressed. In support of this as a possibility, our data demonstrates that RL follicular lymphoma cells express reduced levels of multiple HUSH subunits, suggesting that diminished HUSH activity may directly contribute to the elevated PCDHG expression observed in these cells.

HUSH knockout also upregulates repetitive endogenous DNA elements, including LINE and drives interferon signaling^47,60,64,96,115^, thus it would also be of interest to define if there is a HUSH subunit(s) threshold expression level required to repress PCDHG versus LINE-1 elements. Here it is interesting to note that the EBV transforming program upregulates a range of interferon induced genes, such as IFI44^43^. Thus, there may be a comparatively lower interferon signaling cost for EBV-infected cells to subvert HUSH. In support, TASOR or MORC2 knockout in LCLs did not induce expression of most interferon induced genes.

Burkitt lymphoma cells with the EBV latency I program were not lysed by NK, likely due to insufficient activating NK signals^116,117^. By contrast, EBV transforming programs upregulate CD48 and LFA-1 in newly infected cells and in LCLs^118–121^. While we have not observed changes in HUSH components upon nascent B-cell infection^43^ or EBV reactivation^122^, it will be of interest to determine whether HUSH alters EBV gene regulation in specific infected B or epithelial cell states.

EBV-associated lymphomas including PTLD are increasingly treated by rituximab-based chemotherapy approaches upon detection of elevated EBV viral loads, complicating the opportunity to obtain tissue for transcriptomic, proteomic or spatial analyses. Furthermore, we were unfortunately unable to obtain suitable PCDHG immunostaining signal of formaldehyde fixed human tissues. Therefore, a major future objective will be to develop suitable tools for detection of PCDHG expressions on available formaldehyde fixed paraffin embedded PTLD tissues. Nonetheless, LCLs model key aspects of EBV-transformed B cell states^123^, for instance sharing the latency III program with PTLD tumors. LCLs are also used to develop EBV-specific cytotoxic T cell responses for use in adoptive therapy of PTLD tumors^124–128^.

While our studies focused on NK cells, T cells also play critical roles in control of EBV-infected B cell proliferation^10,129^. An important future objective will be to define the extent to which HUSH inhibition mediated PCDHG expression can blunt T cell responses to EBV infected cells or tumors. Key T cell subsets express NKG2A, including CD8 effector and effector-memory T-cells^130,131^, including in the context of chronic viral infection, in which exhausted CD8+ T cells upregulate NKG2A^132^. Immunodeficiency syndromes further highlight major NKT roles in control of EBV infection^133^, and NKT express NKG2A/CD94.

It would also be of interest to obtain a molecular structure of the PCDHG/NKG2A/CD94 complex, though this is complicated by the propensity of PCDHG to interact *in trans*. Similarly, a future objective will be to systematically define the subset of the 22 PCDHG genes^134^ that bind to and activate NKG2A/CD94 signaling. CD94 can also interact with heparin and sialic acid^135,136^.

Our findings suggest that blockade of PCDHG/NKG2A interaction may be harnessed to re-sensitize particular tumors to NK surveillance. Anti-NKG2A therapies are in development, including the monoclonal antibody monalizumab^131,137^. Indeed, NKG2A blockade by monalizumab enhanced NK killing of EBV-superinfected K562 cells *in vitro*, though in this system, this was likely at the level of NKG2A/HLA-E interaction^138^. We anticipate that antibodies or small molecule therapeutics might be developed to selectively impair PCDHG/NKG2A interaction for use in immunotherapy approaches, including for EBV-driven PTLD. In support, we found that either antibodies against NKG2A or the PCDHG-EC1-3-Fc fusion protein could upregulate NK activity against PCDHG+ lymphoblastoid B-cell targets. Given the increasing interest in developing adoptive transfer NK therapy approaches, it may also be beneficial to engineer NKG2A deficient NK, to minimize the potential for target cell escape through PCDHG de-repression or if elevated PCDHG is detected in the target tissue pre-therapy. These might be particularly relevant to apply to the adoptive NK therapies currently in clinical development for FL^139^.

In conclusion, our CRISPR screen analyses highlighted that HUSH is required to repress PCDHG expression in EBV-transformed lymphoblastoid B cells and to maintain their sensitivity to NK surveillance. In the absence of HUSH activity, PCDHG expressed at the plasma membrane interacts with NKG2A/CD94 complexes to inhibit NK activity. HUSH subunit TASOR KO protected LCL xenografts from adoptively transferred NK cells *in vivo*. EBV uninfected cancers, including FL and neuroblastoma, may subvert PCDHG signaling to evade NK surveillance.

## Resource availability

### Lead Contact

Requests for further information and resources should be directed to and will be fulfilled by the lead contact, Benjamin E. Gewurz.

### Materials availability

All unique/stable reagents generated in this study are available from the lead contact without restriction.

### Data and code availability

All RNA-Seq and CRISPR screen datasets have been deposited in NCBI’s Gene Expression Omnibus (GEO) under PRJNA1458845 and PRJNA1458851.

## Acknowledgements

We thank Dr. Judy Lieberman for sharing the YT-INDY cell line. We thank Drs. Hyung Suk Oh and David Knipe for sharing the SH-SY5Y cell line. We thank Dr. Jose Miguel Lopez-Botet Arbona for sharing the anti-NKG2A Z199 antibody. This work was supported by American Cancer Society Post-doctoral fellowships PF-23-1144614-01-IBCD to Y.S, by NIH award T32AI700245 to J.S.P, by R21AI181873 and R01 CA228700 BEG and by U01CA275301 to BEG and LGR. J.A. is an Investigator of the Howard Hughes Medical Institute. Dr. Gewurz acknowledges generous support from George and Sandra K. Schussel.

## Author contributions

Y.S., E.M.B. and L.Z. performed experiments and data analysis. W.L., J.S.P. and J.A. generated the Fc-fusion PCDHG protein, NKG2A/CD94 soluble proteins, and provided the membrane-associated protein targeting sgRNA library. I.K., H.B., S.S. and L.G.R performed the xenograft experiments. Y.S. and B.E.G. supervised the study. Y.S. and B.E.G. wrote the manuscript. Review and editing of the manuscript were performed by all authors. Y.S. and I.K. contributed equally to this work.

## Materials and Methods

### Cell lines

HEK293T, RL, Reh, Jijoye and NK-92 cells were obtained from American Type Culture Collection (ATCC). GM12878 and GM15892 lymphoblastoid cell lines were purchased from Coriell Institute for Medical Research. P3HR1 cells were from Elliott Kieff. YT-INDY cells and K562 cells were obtained from Judy Lieberman. SH-SY5Y cells were from David Knipe. MUTU I and MUTU III Burkitt cells were obtained from Jeff Sample. HBL1 cells were obtained from Ethel Cesarman. Cell lines with stable Streptococcus pyogenes Cas9 expression were generated by lentiviral transduction and blasticidin selection (5 μg/ml). All B cells and NK cells were grown in RPMI 1640 medium (Gibco, Life Technologies) with 10% fetal bovine serum (FBS, Gibco). HEK293T cells were cultured in Dulbecco’s Modified Eagle’s Medium (DMEM) with 10% FBS. Expi293F^TM^ cells (Thermo Fisher Scientific A14527) were maintained in Expi293 Expression Medium (Thermo Fisher Scientific). All cells were cultured in a humidified incubator at 37°C with 5% CO_2_ and routinely certified as mycoplasma-free, using the MycoAlert kit (Lonza).

### Human Sample

Human tonsil tissue was sourced from Cincinnati Children’s Hospital Medical Center, and leukocyte fractions from platelet donations were obtained from the Brigham and Women’s Hospital Blood Bank. All human materials were discarded, de-identified samples and were expected to include donors of both sexes. Experiments using these cells were performed under non-human subjects research protocols approved by the Mass General Brigham Institutional Review Board (IRB #2022P001270). Informed consent was obtained from all donors by the Brigham and Women’s Hospital Blood Bank and Cincinnati Children’s Hospital Medical Center prior to donation.

### Primary human NK cells

Primary human NK cells were isolated from leukocyte fractions using EasySep Human CD56 Pos Sel Kit II (Stemcell). For activation and expansion, the NK cells were cultured for over one week in the presence of 50 ng/mL interleukin-15 (IL-15, Biolegend) and 5 μM nicotinamide (Sigma)^140^.

### CRISPR/Cas9 loss-of-function screens

To generate libraries for the NK cell cytotoxicity screen, we transduced 130 million Cas9+ GM15892 LCLs with the Broad Institute Brunello sgRNA library^39^ at an MOI of 0.3. Transduction was performed by spinoculation in 12-well plates at 300g for 2 hours in the presence of 4 μg/μl polybrene. Most human genes are targeted by four independent Brunello sgRNAs with unique targeting sequences and PAM sites. Cells were then incubated at 37°C and 5% CO₂ for 6 hours before being pelleted and resuspended in fresh RPMI/FCS medium. Two days later, transduced cells were selected with 3 μg/ml puromycin. The sgRNA-modified B cell libraries were co-cultured with YT-INDY cells or primary NK cells at a 1:1 effector-to-target (E:T) ratio for 7 days. To remove any residual NK cells, blasticidin (100 μg/ml) was added for an additional 7 days.

Surviving B cells were counted and subjected to a second round of NK co-culture and blasticidin treatment. For controls, B cells were maintained in culture without NK cells and passaged every 72 hours over the same time period, keeping at least 40 million cells at each passage to preserve library complexity. After two rounds of selection, genomic DNA was extracted from 40 million cells per replicate using the Qiagen Blood and Cell Culture DNA Maxi Kit according to the manufacturer’s instructions. 2 replicates were performed for both YT-INDY and primary NK screens.

To attach sequencing adapters and barcode each sample, sgDNA was amplified by PCR in multiple 100 μL reactions, each containing up to 10 μg genomic DNA, following the protocol of Doench et al^40^. For each 96-well PCR plate, a master mix was prepared consisting of 75 μL ExTaq DNA Polymerase (Clontech), 1000 μL 10× ExTaq buffer, 800 μL dNTP mix, 50 μL P5 stagger primer mix (100 μM), and 2075 μL water. Each well contained 50 μL sgDNA plus water, 40 μL of the PCR master mix, and 10 μL of a uniquely barcoded P7 primer (5 μM). The PCR conditions were: 95°C for 1 minute; then 28 cycles of 94°C for 30 seconds, 52.5°C for 30 seconds, and 72°C for 30 seconds; followed by a final extension at 72°C for 10 minutes. P5 and P7 primers were synthesized by Integrated DNA Technologies (IDT). PCR products were purified using Agencourt AMPure XP SPRI beads (Beckman Coulter) according to the manufacturer’s protocol. Libraries were sequenced on an Illumina HiSeq2000 platform. Reads were processed by first identifying the CACCG sequence—which is present 5′ to all sgRNA inserts in the vector—in the primary read file. The 20 nucleotides immediately following CACCG were extracted as the sgRNA insert and mapped to a reference of all sgRNAs in the library.

Sample identification was achieved using the 8-nt barcode in the P7 primer to assign each read to its corresponding PCR well or experimental condition. The resulting count matrix was normalized to reads per million (RPM) using the formula: (reads per sgRNA/total reads per condition) × 10⁶. To enable log2 transformation—and to account for sgRNAs with zero reads—one was added to all RPM values prior to log2 transformation.

For the NK receptor screen, we used a lentiviral sgRNA library that targets membrane associated proteins^71,72^ to generate YT-INDY cell libraries following an analogous protocol. This library targets most human transmembrane protein encoding genes with 10 unique sgRNAs.

Following puromycin selection, YT-INDY cells were stained with 16μg/ml PE-conjugated Fc-PCDHGB2-EC1-3 at 37°C for 3 hours. The 5% of cells with the lowest PE staining were isolated by flow cytometry. As a control, 24 million puromycin-selected cells were harvested for genomic DNA extraction. 3 replicates of the screen were performed. SgRNAs were PCR amplified from purified genomic DNA, and the amplified sgRNA sequences were quantified by Illumina HiSeq sequencing. Hit gene identification and statistical significance were determined using the STARS algorithm^40^, applying a stringent threshold of q < 0.05 (FDR-adjusted p-value). For each gene classified as a hit, at least two independent sgRNAs showed significant enrichment.

STARS is a gene ranking algorithm specifically designed for CRISPR-based genetic perturbation screens, and we applied it to assess the rank and statistical significance of screen hits. This approach takes advantage of the abundance data from four independent sgRNAs targeting each gene and compares their performance across the two screen conditions. To ensure specificity, we excluded sgRNAs with more than five predicted genome-wide off-target sites based on the latest datasets from John Doench and the Broad Institute. Most sgRNAs in the Brunello library do not have predicted off-target effects, and the unique off-target signatures for each sgRNA targeting a given gene increase confidence when multiple sgRNAs yield consistent phenotypes. For each sgRNA, we calculated the log2-fold change by comparing data between the control and NK co-cultured cells. Each sgRNA was then ranked by its log2-fold change, and this ranking was normalized by the total number of sgRNAs to yield a percent rank. For STARS analysis, percent-rank values were combined across subpools.

### Antibodies

For immunoblot analysis, antibodies against TASOR (Cell Signaling Technologies, E6M2Y, #92278), PPHLN1 (Invitrogen, #PA5-58583), MPHOSPH8 (Santa Cruz, C-8, #sc-398598), MORC2 (Cell Signaling Technologies, #77667), Protocadherin Gamma (Invitrogen, S159-5, # MA5-27615), DDX1 (Bethyl, #A300-521A), GFP tag (Proteintech, #50430-2-AP), HA tag (Cell Signaling Technologies, C29F4, #3724), and Strep Tag II (Invitrogen, 1810CT579.47.56.10, MA5-37747) were used at 1:1000. Anti-GAPDH (EMD Millipore #MAB374) was used at 1:5000. Cell Signaling Technology HRP-linked anti-mouse (#7076) and anti-rabbit (#7074) antibodies were used as secondary antibodies for immunoblot analysis at 1:5000. Protocadherin Gamma (Invitrogen, S159-5, MA5-27615) was used for immunofluorescence assay at 1:1000. Antibodies targeting FAM208A (TASOR, Atlas Antibodies, HPA006735), SETDB1 (Proteintech, 11231-1-AP), and H3K9me3 (Active Motif, pAb, 39062) were used for ChIP assay. For flow cytometry analysis, Biolegend antibodies targeting CD56 (MEM-188, 362504), CD48 (BJ40, #336714), CD11a (HI111, #301208), IFNγ (4S.B3, 502523), CD107a (H4A3, #328605), IgG Fc (M1310G05, #410707), CD159a (S19004C, #375107) were used. Anti-NKG2A (Z-199) from Dr.

Jose Miguel Lopez-Botet Arbona was used in blocking assays at 10 μg/mL.

### CRISPR/Cas9 mutagenesis studies

For cell lines stably expressing Cas9, sgRNA sequences from the Broad Institute’s Brunello library were utilized. These sgRNA oligonucleotides were inserted into the pLentiGuide-Puro vector (Addgene plasmid #52963, provided by Feng Zhang, Broad Institute of MIT and Harvard), and lentiviral particles were produced in HEK293T cells. Cells underwent two rounds of transduction at 48 and 72 hours after plasmid transfection, followed by selection with 3 μg/mL puromycin for at least four days.

Genome editing in primary B cells was performed as previously described^141^. In brief, Cas9 ribonucleoprotein complexes were delivered via electroporation. Specifically, crRNAs targeting MORC2 were selected using the Alt-R Predesigned Cas9 crRNA Selection Tool (Integrated DNA Technologies). Both tracrRNA and Cas9 Nuclease V3 were obtained from Integrated DNA Technologies as well. crRNA and tracrRNA were annealed to form duplexes and then combined with Cas9 protein for 20 minutes to assemble the RNP complexes. B cells were then electroporated with these RNPs using the Neon NxT Electroporation System (1700 V, 20 ms, 2 pulses). The sgRNA sequences used are listed in **Table S1**.

### Flow cytometry

For surface marker staining, 1 × 10^6^ cells were washed twice in FACS buffer (PBS supplemented with 1% FBS) before incubation with fluorochrome-conjugated antibodies against surface proteins for 30 minutes in the dark. For intracellular staining, cells were first fixed using eBioscience™ IC Fixation Buffer immediately after surface staining, then washed twice with eBioscience™ Permeabilization Buffer, followed by a 30-minute incubation with antibodies against intracellular targets in the dark. To discriminate live and dead cells, samples were incubated with 7-AAD, propidium iodide (PI), or Zombie dye (Biolegend) for 10 minutes on ice. After all staining steps, cells were washed with FACS buffer and analyzed by flow cytometry.

Data was processed using FlowJo X software (FlowJo). For cell viability staining, Biolegend Zombie NIR™ Fixable Viability Kit or Zombie Aqua™ Fixable Viability Kit were used following manufacture’s protocol.

### In vitro NK cytotoxicity assay

To evaluate the cytotoxic activity of NK cells, an *in vitro* cytotoxicity assay was conducted. Briefly, target cells were labeled with Invitrogen CFSE, CellTrace Violet (CTV) or CellTrace Far Red (CTFR) as fluorescent tracking dyes according to the manufacturer’s protocol. Labeled target cells were then co-cultured with NK cells at the indicated effector-to-target (E:T) ratios for defined time periods, typically 24 hours, at 37°C. After incubation, cells were harvested and stained with viability dyes to discriminate live and dead cells. Flow cytometry was used to determine the proportions and absolute counts of live CFSE^+^ (or CTV^+^) target cells among the total live population. The cytotoxicity (killing efficiency) was calculated by comparing the number of live CFSE^+^ target cells remaining after NK cell co-culture to the number of live CFSE^+^ cells in control wells cultured without NK cells for the same duration.

### In vitro adhesion assay

To assess the adhesion between NK cells and target cells, an *in vitro* adhesion assay was carried out as previously described^142^. Briefly, NK cells were labeled with CellTrace Violet, while target cells were stained with CFSE as a distinguishing marker. The two cell populations were mixed at a 1:1 ratio and incubated at 37°C for 0, 30, or 120 minutes. Following incubation, the cell mixtures were fixed in 2% paraformaldehyde (PFA) in PBS. The extent of cell adhesion was then quantified by flow cytometry, identifying double-positive (CFSE and CellTrace Violet) events.

### Apoptosis assays

For induction of apoptosis, cells were treated with 50 ng/mL tumor necrosis factor alpha (TNFα; Sigma) and 10 μg/mL cycloheximide (CHX; R&D Systems). To induce necroptosis, cells were exposed to 50 ng/mL TNFα, 10 μg/mL CHX, and 20 μM z-VAD-FMK (Apexbio). All treatments were performed for the indicated durations under standard culture conditions.

### Immunoblot analysis

Cells were harvested and normalized using the CellTiterGlo (CTG) luciferase assay (Promega, Cat#G7570). Following normalization, cell pellets were lysed in 1x Laemmli Sample Buffer (Bio-Rad) with ultrasonic disruption, then heated at 95 °C for 10 minutes. The resulting lysates were separated by SDS-PAGE and transferred to nitrocellulose membranes (Bio-Rad). Membranes were blocked in 5% nonfat dry milk in TBST for 1 hour, then incubated overnight at 4 °C with primary antibodies. After washing three times with TBST, membranes were incubated with secondary antibodies for 1 hour at room temperature. Following three additional washes with TBST, blots were developed with ECL reagent (Thermo Fisher #34578) and visualized using the Licor Fc imaging system and the Image Studio Lite 5.2 software (LICORbio).

### Proliferation assay

Cells were plated at a density of 0.5 million cells per well in 12-well plates. The culture medium was refreshed every 3 to 4 days, and viable cell numbers were determined on specified days using trypan blue exclusion. Cell counts were normalized to those on day 0 and plotted accordingly.

### RNAseq analysis

Total RNA was isolated from cells using the RNeasy Mini Kit (Qiagen) according to the manufacturer’s protocol, including an on-column DNase treatment to remove residual genomic DNA. To prepare libraries, polyadenylated RNA was purified from 1 μg of DNA-free total RNA using the NEBNext Poly(A) mRNA Magnetic Isolation Module (New England Biolabs), and libraries were generated with the NEBNext Ultra RNA Library Prep Kit (New England Biolabs). Library integrity and size distribution were evaluated using an Agilent Bioanalyzer with a DNA chip. Indexed libraries were pooled and sequenced on an Illumina NextSeq 500 instrument, producing single-end 75 base pair reads at the Dana Farber Molecular Biology core facility. Raw gene expression data were quantified using salmon (version 1.2.0)^143^ mapped to the human GENCODE v28 (GRCh37) gene annotation, and subsequent differential gene expression analysis was performed using DESeq2^144^. For heatmap analysis, Z-scores of normalized mRNA reads are shown. The Z-score is calculated by subtracting the mean from an individual data point and dividing the result by the standard deviation.

### ChIP-qPCR

Twenty million B cells were harvested and cross-linked by incubating with 1% formaldehyde for 10 minutes at room temperature. Cross-linking was halted by treating the cells with 0.3 M glycine for an additional 5 minutes, followed by two washes with ice-cold PBS. Cells were subsequently lysed in ChIP lysis buffer containing 1% SDS, 10 mM Tris-HCl (pH 8.0), 10 mM EDTA, and 1× cOmplete™ EDTA-free protease inhibitor cocktail (Sigma) on ice for 20 minutes. Chromatin was sheared by sonication to generate DNA fragments of the desired size. After debris removal by centrifugation, the clarified chromatin was diluted with ChIP dilution buffer (16.7 mM Tris-HCl pH 8.0, 1.2 mM EDTA, 167 mM NaCl, 1.1% Triton X-100, 0.01% SDS, supplemented with protease inhibitors). The diluted chromatin was then incubated overnight at 4°C with 4 μg of either target-specific antibody or control IgG, along with pre-washed protein A magnetic beads (Thermo Fisher), under gentle rotation. The immune complexes were washed sequentially using a series of buffers: twice with low salt buffer (20 mM Tris-HCl pH 8.0, 2 mM EDTA, 150 mM NaCl, 1% Triton X-100, 0.1% SDS), twice with high salt buffer (20 mM Tris-HCl pH 8.0, 2 mM EDTA, 500 mM NaCl, 1% Triton X-100, 0.1% SDS), once with LiCl buffer (10 mM Tris-HCl pH 8.0, 1 mM EDTA, 0.25 M LiCl, 1% NP-40, 1% sodium deoxycholate), and once with TE buffer (10 mM Tris-HCl pH 8.0, 1 mM EDTA). To elute immunoprecipitated chromatin and reverse the cross-links, beads were resuspended in elution buffer (100 mM NaHCO₃, 1% SDS) and incubated at 65°C for 2 hours in the presence of Proteinase K. Purified DNA was obtained using the QIAquick PCR purification kit (Qiagen). Quantification of immunoprecipitated DNA was performed by quantitative PCR and normalized to input controls. Primers used for ChIP-qPCR were listed in **Table S2**.

### Recombinant protein production

We cloned human PCDHGB2 EC1-3 (M1-V105 including the native signal peptide sequence, GenBank NM_018923.3) into the pVRC expression vector, which encodes human IgG1 Fc as a C-terminal fusion protein. An HA tag (YPYDVPDYA) and a human rhinovirus 3C protease cleavage site (LEVLFQGP) were inserted between EC3 and the C-terminal Fc, with an AS or GS linker flanking the insertion. The pVRC-PCDHG-EC1-3 plasmid was transfected into Expi 293F cells (Thermo Fisher A14527) using the ExpiFectamineTM 293 Transfection Kit (Thermo Fisher A14525) according to the manufacturer’s protocol. At 5 days post-transfection, we purified the Fc-fusion protein using MabSelect SuRe LX protein A affinity resin (GE Healthcare 17549802), according to the manufacturer’s protocol. Proteins were stored in a high-salt buffer (20 mM Tris, 1M NaCl, pH 7.5).

We cloned the human NKG2A (KLRC1) extracellular segment (Q99–L233, GenBank X54867.1) into the pVRC expression vector downstream of a tissue plasminogen activator (TPA) signal sequence (MKRGLCCVLLLCGAVFVSPS), a GS linker, a twin-strep tag (WSHPQFEKGGGSGGGGSGGSAWSHPFEK), an AS linker, a human rhinovirus 3C protease cleavage site, and a second GS linker. We cloned the human CD94 (KLRD1) extracellular segment (F35-I139, GenBank U30610.1) into the pVRC expression vector downstream of a TPA signal sequence, a GS linker, a 6xHis tag (HHHHHH), an AS linker, a human rhinovirus 3C protease cleavage site, and a second GS linker. To express the heterodimer of the extracellular segments of NKG2A and CD94, we transfected pVRC-twin-strep-NKG2A and pVRC-6xHis-CD94 into Expi 293F cells (Thermo Fisher A14527) at a 1:1 ratio using the ExpiFectamineTM 293 Transfection kit (Thermo Fisher A14525). Five days post-transfection, we purified the soluble NKG2A-CD94 complex from culture supernatant using the Strep-Tactin® XT Sepharose chromatography resin (Cytiva 29401324), according to the manufacturer’s protocol. The eluted protein was further purified on size-exclusion chromatography using a Superdex 200 increase column. The protein was stored in TBS buffer (20 mM Tris, 150 mM NaCl, pH 7.5).

### Fc-Fusion Protein Binding Assay

Cells were incubated with the Fc-fusion protein, which was diluted in H₂O to the indicated concentrations, for 3 hours at 37 °C. After incubation, cells were washed to remove unbound protein and stained with PE anti-human IgG Fc antibody (M1310G05, BioLegend) prior to flow cytometric analysis. For NK-92 cells, Human TruStain FcX™ (BioLegend) was applied according to the manufacturer’s instructions before protein incubation.

### In vitro binding assay

Fc-PCDHGB2-EC1-3 and Strep-KLRC1/his-KLRD1 were mixed at 1:1 ratio in Co-IP lysis buffer (1% v/v NP40, 150 mM Tris, 300 mM NaCl, 1 X cOmpleteTM EDTA-free protease inhibitor cocktail (Sigma), 1 mM Na_3_VO_4_ (Sigma) and 1 mM NaF (Sigma)) and incubated at 4°C with gentle rotation for 4 hours. 10% (v/v) of an aliquot was preserved as the input. The rest were incubated with 20 uL magnetic HA beads (Thermo Fisher) with gentle rotation for overnight. Following multiple washes with the Co-IP lysis buffer, the beads were collected using a magnet and boiled at 95 °C for 10 mins in 2x Laemmli Sample Buffer (Biorad). The supernatant was subjected to immunoblot analysis.

### Immunofluorescence assay

Confocal microscopy was performed as previously described^145^. Cells were seeded onto 12-well chamber slides (ibidi) and cultured for 24 hours. Following 2 times of PBS washes, slides were fixed in 4% paraformaldehyde for 10 minutes, then permeabilized with 0.5% Triton X-100 in PBS and blocked with 1% BSA in PBS. For immunostaining, slides were incubated with primary antibody in a humidified chamber at 37°C for 4 hours, followed by secondary antibody incubation at 37°C for 1 hour. After staining, cells were washed twice with PBS and mounted overnight using ProLong™ Gold Antifade Mountant with DAPI. Confocal imaging and analysis were performed using a Zeiss LSM 800 microscope and Zeiss Zen Lite (Blue) software.

### Mouse xenograft experiments

Murine xenograft experiments were conducted in accordance with Institutional Animal Care & Use Committee (IACUC #2022-0002) protocol regulations at Weill Cornell Medical Center.

NOD.Cg-Prkdcscid Il2rgtm1Wjl Tg(IL15)1Sz/SzJ (NSG-IL15) mice, aged 6-8 weeks, were purchased from Jackson Laboratories. Sixteen mice per group were injected subcutaneously in each flank with 1×10^7^ wild-type (WT) or TASOR knockout (KO) lymphoblastoid cell lines (LCLs) suspended in PBS with Matrigel. Once tumors reached approximately 500mm3, mice were further subdivided to receive intravenous injections of either the NK92 natural killer cell lines or

PBS control. Tumor growth was monitored at least twice weekly using digital calipers, and tumor volume was calculated using the formula V = (LxWxW)/2. Statistical significance for tumor growth over time was assessed by two-way ANOVA. Additionally, tumor growth was quantified by calculating the area under the curve (AUC) using the pracma package in RStudio, with statistical significance determined by multiple t-test. Data normality was assessed using Shapiro-wilk normality test. Mice were euthanized upon reaching the experimental endpoint, defined by tumor size. Tumors were harvested for flow cytometry. NK cell infiltration into tumors was assessed by flow cytometry.

### Tonsil organoids model

Tonsil organoids were prepared as previously described^76^. In brief, tonsil tissue was minced and filtered through a 100 μm nylon strainer to obtain a single cell suspension. Tonsil cells were cryopreserved in FBS with 10% DMSO at −135°C until use. CD19^+^ B cells were isolated by magnetic bead negative selection from human tonsil cell suspensions, and electroporated with Cas9 RNP loaded with nontargeting control or MORC2 sgRNA. Control cells were stained with 2mM Celltrace Violet (CTV), MORC2 KO cells were stained with 0.5mM Celltrace Far Red (CTFR). 0.2 million of these cells (mixed at a 1:1 ratio) were cultured alone or together with 6 million autologous tonsil cells in transwell plate tonsil organoid cultures. The organoid media was composed of RPMI1640 with 1x glutamax (Gibco), 10% FBS, 1x non-essential amino acids (Gibco), 1x sodium pyruvate (Gibco), 1x penicillin/streptomycin, 1x Normocin (InvivoGen), 1x insulin/selenium/transferrin cocktail (Gibco) and 1 μg/mL of human BAFF (Biolegend). Cells were infected with B95.8 EBV at an MOI of 0.1. On day 5 post infection, the %CTV+ and CTFR+ cells were analyzed by FACS.

### Statistical analysis

Statistical significance between groups was determined using either an unpaired Student’s t-test or ANOVA followed by suitable post-hoc tests, performed in GraphPad Prism 9 software. P values were represented as follows: ns = not significant, p > 0.05; *p < 0.05; **p < 0.01; ***p < 0.001; ****p < 0.0001. Figures were drawn with commercially available GraphPad, BioRender, and Microsoft PowerPoint software.

## Supplemental Figure Legends

**Figure S1. CRISPR Screen for LCL Factors that Support NK Surveillance, Related to Figure 1**.

A) YT-INDY killing of GM15892 LCLs. FACS analysis of GM15892 cell vital dye 7-AAD uptake following five days of co-incubation with YT-INDY NK cells at an effector to target ratio of 1:1. Representative of n=3 replicates.

B) Primary human NK killing of GM15892 LCLs. FACS analysis of GM15892 cell vital dye 7-AAD uptake following five days of co-incubation with primary human peripheral blood IL-15/NAM treated NK cells at an effector to target ratio of 1:1. Representative of n=3 replicates.

C) Rug plot analysis of the indicated screen hit sgRNAs (pink) versus other Brunello library sgRNAs (gray) from the YT-INDY CRISPR screen.

D) Rug plot analysis of the indicated screen hit sgRNAs (pink) versus other Brunello library sgRNAs (gray) from the primary human NK CRISPR screen.

**Figure S2. Screen hit validation, related to Figure 1**.

A) Immunoblot analysis of whole cell lysates (WCL) from Cas9+ GM15892 LCLs that expressed screen the indicated control versus screen hit sgRNAs. Blots are representative of n=3 replicates.

B) FACS analysis of plasma membrane CD48 (left) versus ITGAL-encoded LFA-1 subunit CD11A (right) on Cas9+ GM15892 LCLs that expressed the indicated sgRNA. Data are representative of n=3 replicates.

C) Growth curve analysis of Cas9+ GM15892 cells that expressed the indicated sgRNAs. Shown are mean ± standard deviation (SD) live cell values from n=3 independent replicates.

Statistical significance was assessed by two-tailed unpaired Student’s t test (C). ns, not significant.

**Figure S3. HUSH KO effects on target cell killing, related to** Figure 2.

A) HUSH KO effects on GM12878 LCL killing by NK. Mean + SD relative live cell numbers from n=3 replicates of Cas9+ GM12878 that expressed the indicated control (Ctrl), TASOR or MORC2 sgRNAs and co-cultured at a 2:1 ratio with YT-INDY (top), NK-92 (middle) or primary human IL-15/NAM treated NK (bottom) for 24 hours. Numbers were normalized to LCLs cultured in the absence of NK. Shown at bottom are representative immunoblots of WCL from GM12878 that expressed the indicated sgRNAs. B) TASOR KO effects on B cell lymphoma or K562 killing by NK. Mean + SD relative live cell numbers from n=3 replicates of Cas9+ cells that expressed the indicated control (white bar) or TASOR (orange bar) sgRNAs and co-cultured at a 2:1 ratio with YT-INDY (top) or NK-92 (bottom) for 24 hours. Numbers were normalized to cells cultured in the absence of NK. Representative immunoblots of WCL from cells expressing control or TASOR sgRNAs are shown at bottom. Blots run separately are indicated by the borders.

Statistical significance was assessed by one-way ANOVA followed by Tukey’s multiple comparisons test (A) or two-tailed unpaired Student’s t test (B). ns, not significant, *P < 0.05, **P < 0.01, ***P < 0.001, ****P < 0.0001.

**Figure S4. LCL HUSH KO effects on NK adherence, Related to** Figure 2.

A) Effects of LCL HUSH NK on primary human NK adherence. Shown are representative FACS plots of *in vitro* adhesion assays, in which IL-15/NAM-activated primary human NK cells were co-cultured with Cas9+ GM15892 LCLs for the indicated minutes. NK were stained with CellTrace Violet (CTV) and GM15892 LCLs were stained with carboxyfluorescein succinimidyl ester (CFSE) just prior to the co-incubation.

B) Mean + SD percentages of adherent GM15892/primary human NK cell pairs from n=3 independent replicates as in (A).

C) Effects of LCL HUSH NK on YT-INDY GM15892 NK adherence. Shown are representative FACS plots of *in vitro* adhesion assays, in which YT-INDY NK cells were co-cultured with Cas9+ GM15892 LCLs for the indicated minutes. NK were stained with CTV and GM15892 LCLs were stained with CFSE just prior to the co-incubation. Mean + SD percentages of adherent GM15892/YT-INDY NK cell pairs from n=3 independent replicates.

D) Effects of LCL HUSH NK on GM12878 LCL YT-INDY NK adherence. Shown are representative FACS plots of *in vitro* adhesion assays, in which YT-INDY NK cells were co-cultured with Cas9+ GM12878 LCLs for the indicated minutes. NK were stained with CTV and GM12878 LCLs were stained with CFSE just prior to the co-incubation. Mean + SD percentages of adherent GM12878/YT-INDY NK cell pairs from n=3 independent replicates.

Statistical significance was assessed by one-way ANOVA followed by Tukey’s multiple comparisons test (B-D). ns, not significant, *P < 0.05, **P < 0.01, ***P < 0.001, ****P < 0.0001.

**Figure S5. HUSH KO effects on NK activation. Related to** Figure 2.

A) GM15892 LCL HUSH KO effects on YT-IDY IFNγ expression and plasma membrane CD107a. Mean + SD values from n=3 replicates of IFNγ+ (left) and CD107a+ (right) of YT-INDY cells that were cultured for 5 hours alone or with Cas9+ GM15892 LCLs that expressed the indicated sgRNAs. Cultures were treated with protein transport inhibitor for 5 hours prior to IFNγ analysis.

B) GM12878 LCL HUSH KO effects on YT-IDY IFNγ expression and plasma membrane CD107a. Mean + SD values from n=3 replicates of IFNγ+ (left) and CD107a+ (right) of YT-INDY cells that were cultured for 5 hours alone or with Cas9+ GM12878 LCLs that expressed the indicated sgRNAs. Cultures were treated with protein transport inhibitor for 5 hours prior to IFNγ analysis.

C) Analysis of HUSH KO effects on apoptosis and necroptosis induction. Shown are Mean + SD values of %7-AAD+ (dead) cells of Cas9+ GM15892 LCLs that expressed the indicated sgNRA and that were treated for 24 hours with TNFα (50ng/mL) and cycloheximide (CHX, 10μg/mL) for induction of apoptosis or with TNFα, CHX and caspase-inhibitor zVAD-Fmk (20μM) for induction of necroptosis.

D) Effects of exposure to HUSH KO LCL on subsequent NK killing. NK were pre-incubated with Cas9+ GM12878 with the indicated sgRNA for 4 days, and then used for killing assay with fresh control LCL targets. Mean + SD live cell numbers from n=3 replicates of GM12878 following co-culture with YT-INDY (left) or NK-92 (right) at a 2:1 E:T ratio for 24 hours. Values were normalized by live cell values of GM12878 cultured in the absence of NK.

Statistical significance was assessed by one-way ANOVA followed by Tukey’s multiple comparisons test (A-D). ns, not significant, *P < 0.05, **P < 0.01, ***P < 0.001, ****P < 0.0001.

**Figure S6. HUSH KO effects on EBV and endogenous retroviral or LINE expression, related to** Figure 3.

A) Analysis of TASOR (left) or MORC2 (right) sgRNA effects on EBV gene expression.

Shown are volcano plot analyses from n=3 RNAseq replicates of GM15892 that expressed control versus TASOR sgRNAs (left panel) or control versus MORC2 sgRNAs (right panel). Selected EBV genes are highlighted in red circles.

B) Heatmap analysis of the expression of human genome endogenous retroviral or LINE/L1 elements in GM15892 LCLs with control, TASOR or MORC2 sgRNAs from RNAseq analysis. Shown are the endogenous human genome elements whose expression was most highly changed by TASOR or MORC2 depletion.

**Figure S7. Analysis of the LCL PCDHGA1 locus, related to** Figure 3.

A) Immunoblot analysis of WCL from GM12878 LCLs that expressed the indicated sgRNAs. Blots are representative of n=3 experiments.

B) H3K9me3 ChIP-seq tracks at the GM12878 LCL *PCDHG* locus. Data was obtained from the ENCODE project (accession: ENCSR000AOX).

C-E) ChIP-qPCR analysis of the GM15892 LCL *PCDHGA2* locus. Mean + SD values from n=3 ChIP-qPCR replicates of TASOR (C) or SETDB1 (D) occupancy or of H3K9me3 (E) levels at the *PCDHGA2* promoter region. Anti-TASOR, SETDB1 and H3K9me3 ChIP were performed on chromatin from GM15892 LCLs that expressed the indicated control or TASOR sgRNAs, followed by qPCR.

Statistical significance was assessed by two-tailed unpaired Student’s t test (C-E).

**Figure S8. Analysis of PCDHG expression effects on NK-mediated LCL lysis, related to** Fig. 4.

A) Immunoblot analysis of WCL from GM12878 LCLs that expressed the indicated control or *PCDHG* common exon targeting sgRNAs, and that also expressed control, TASOR or MORC2 sgRNAs, as indicated.

B) Analysis of PCDHG knockout on HUSH KO resistance to YT-INDY mediated lysis.

Mean + SD relative live cell values from n=3 replicates of Cas9+ GM12878 LCLs with the indicated sgRNAs following co-culture with YT-INDY at an E:T of 2:1 or culture alone for 24 hours.

C) Analysis of PCDHG knockout on HUSH KO resistance to NK92 mediated lysis.

Mean + SD relative live cell values from n=3 replicates of Cas9+ GM12878 LCLs with the indicated sgRNAs following co-culture with NK-92 at an E:T of 2:1 or culture alone for 24 hours.

D) Analysis of PCDHGB2 expression effects on LCL protection from YT-INDY lysis.

Mean + SD relative live cell numbers from n=3 replicates of GM12878 LCL that were electroporated with empty vector versus PCDGHB2 vectors and that were then co-cultured with YT-INDY at a 2:1 E:T ratio or alone for 24 hours. YT-INDY co-cultured LCL live cell values were normalized by values from identical LCLs cultured alone.

I) E) Analysis of PCDHGB2 expression effects on LCL protection from NK-92 lysis. Mean + SD relative live cell numbers from n=3 replicates of GM12878 LCL that were electroporated with empty vector versus PCDGHB2 vectors and that were then co-cultured with NK-92 at a 2:1 E:T ratio or alone for 24 hours. NK-92 co-cultured LCL live cell values were normalized by values from identical LCLs cultured alone.

F) Analysis of LCL PCDHGB2 transgene expression. Shown are representative confocal microscopy images of wildtype or truncation mutant PCDHGB2 cDNA expression in GM15892 LCLs. Cells were transduced with lentiviral expression vectors and puromycin selected prior to immunostaining.

G) Analysis of recombinant Fc-PCDHGB2-EC1-3. Representative Coomassie blue stained gel loaded with purified Fc-PCDHGB2-EC1-3 loaded in non-reducing versus reducing loading buffer with 5% β-mercaptoethanol.

H) Analysis of Fc-PCDHGB2-EC1-3 labeling of HEK-293 plasma membranes. Shown are Mean ± SD MFI values from n=3 replicates of FACS analysis of control IgG versus Fc-PCDHGB2-EC1-3 stained HEK-293. Cells were treated with the indicated IgG versus Fc-PCDHGB2-EC1-3 concentrations and then stained with PE-tagged anti-IgG Fc prior to FACS analysis.

I) Analysis of Fc-PCDHGB2-EC1-3 labeling of NK-92 plasma membranes. Shown are Mean ± SD MFI values from n=3 replicates of FACS analysis of control IgG versus Fc-PCDHGB2-EC1-3 stained NK-92 cells. Cells were pretreated with Fc receptor blocking solution for 15 minutes, then treated with the indicated IgG versus Fc-PCDHGB2-EC1-3 concentrations and stained with PE-tagged anti-IgG Fc prior to FACS analysis.

J) Analysis of Fc-PCDHGB2-EC1-3 effects on NK-92 mediated LCL killing. Mean + SD relative live cell numbers from n=3 replicates of GM15892 cultured alone or with NK-92 that were pre-treated with 16 IgG or with Fc-PCDHGB2-EC1-3 μg/mL for 24 hours. LCLs and NK-92 were co-cultured at a 2:1 E:T ratio for 24 hours.

Statistical significance was assessed by two-tailed unpaired Student’s t test (D,E,J) or two-way ANOVA followed by Tukey’s multiple comparisons test (B,C,H,I). *P < 0.05, **P

< 0.01, ***P<0.001, ****P < 0.0001, ns, not significant.

**Figure S9. Analysis of CRISPR depletion of KLRC1/NKG2A, related to** Figure 5.

A) Representative FACS analysis of plasma membrane NKG2A expression on Cas9+ YT-INDY cells that expressed control or independent screen hit KLRC1 sgRNAs.

B) Representative FACS analysis of plasma membrane NKG2A expression on Cas9+ NK-92 cells that expressed control or KLRC1 sgRNAs.

C) Size-exclusion chromatogram (SEC) elution traces of recombinant CD94/NKG2A.

**Figure S10. Schematic model of key HUSH roles in PCDHG expression and in EBV-transformed lymphoblastoid B cell NK surveillance.**

EBV transformed B cells highly express CD48, which stimulates NK cell activation and infected B-cell lysis. The HUSH complex represses PCDHG expression to support EBV-transformed B cell driven NK activation. In the absence of HUSH activity, PCDHG is de-repressed, traffics to the plasma membrane and associates with NKG2A/CD94 to deliver an inhibitory signal that overcomes CD48/2B4 NK activation. This serves to support infected B cell escape from NK surveillance.

**Table S1. sgRNA or crRNA sequences used in this study**

**Table S2. ChIP-qPCR primer sequences used in this study**

## References

1 Farrell, P. J. Epstein-Barr Virus and Cancer. Annu Rev Pathol 14, 29–53, doi:10.1146/annurev-pathmechdis-012418-013023 (2019).

2 Munz, C. Latency and lytic replication in Epstein-Barr virus-associated oncogenesis. Nat Rev Microbiol 17, 691–700, doi:10.1038/s41579-019-0249-7 (2019).

3 Cohen, J. I., Mocarski, E. S., Raab-Traub, N., Corey, L. & Nabel, G. J. The need and challenges for development of an Epstein-Barr virus vaccine. Vaccine 31 Suppl 2, B194–196, doi:10.1016/j.vaccine.2012.09.041 (2013).

4 Damania, B., Kenney, S. C. & Raab-Traub, N. Epstein-Barr virus: Biology and clinical disease. Cell 185, 3652–3670, doi:10.1016/j.cell.2022.08.026 (2022).

5 Chiu, Y. F., Ponlachantra, K. & Sugden, B. How Epstein Barr Virus Causes Lymphomas. Viruses 16, doi:10.3390/v16111744 (2024).

6 Parvaneh, N., Filipovich, A. H. & Borkhardt, A. Primary immunodeficiencies predisposed to Epstein-Barr virus-driven haematological diseases. Br J Haematol 162, 573–586, doi:10.1111/bjh.12422 (2013).

7 Cohen, J. I. Primary Immunodeficiencies Associated with EBV Disease. Curr Top Microbiol Immunol 390, 241–265, doi:10.1007/978-3-319-22822-8_10 (2015).

8 Tangye, S. G. & Latour, S. Primary immunodeficiencies reveal the molecular requirements for effective host defense against EBV infection. Blood 135, 644–655, doi:10.1182/blood.2019000928 (2020).

9 Lurain, K. A. et al. HIV-associated cancers and lymphoproliferative disorders caused by Kaposi sarcoma herpesvirus and Epstein-Barr virus. Clin Microbiol Rev 37, e0002223, doi:10.1128/cmr.00022-23 (2024).

10 Taylor, G. S., Long, H. M., Brooks, J. M., Rickinson, A. B. & Hislop, A. D. The immunology of Epstein-Barr virus-induced disease. Annu Rev Immunol 33, 787–821, doi:10.1146/annurev-immunol-032414-112326 (2015).

11 Desimio, M. G., Covino, D. A., Rivalta, B., Cancrini, C. & Doria, M. The Role of NK Cells in EBV Infection and Related Diseases: Current Understanding and Hints for Novel Therapies. Cancers (Basel) 15, doi:10.3390/cancers15061914 (2023).

12 Chijioke, O., Landtwing, V. & Munz, C. NK Cell Influence on the Outcome of Primary Epstein-Barr Virus Infection. Front Immunol 7, 323, doi:10.3389/fimmu.2016.00323 (2016).

13 Png, Y. T., Yang, A. Z. Y., Lee, M. Y., Chua, M. J. M. & Lim, C. M. The Role of NK Cells in EBV Infection and EBV-Associated NPC. Viruses 13, doi:10.3390/v13020300 (2021).

14 Mace, E. M. & Orange, J. S. Emerging insights into human health and NK cell biology from the study of NK cell deficiencies. Immunol Rev 287, 202–225, doi:10.1111/imr.12725 (2019).

15 Fleisher, G. et al. A non-x-linked syndrome with susceptibility to severe Epstein-Barr virus infections. J Pediatr 100, 727–730, doi:10.1016/s0022-3476(82)80572-6 (1982).

16 Mace, E. M. et al. Biallelic mutations in IRF8 impair human NK cell maturation and function. J Clin Invest 127, 306–320, doi:10.1172/JCI86276 (2017).

17 Nakid-Cordero, C., Baron, M., Guihot, A. & Vieillard, V. Natural Killer Cells in Post-Transplant Lymphoproliferative Disorders. Cancers (Basel) 13, doi:10.3390/cancers13081836 (2021).

18 Flórez-Álvarez, L., Hernandez, J. C. & Zapata, W. NK Cells in HIV-1 Infection: From Basic Science to Vaccine Strategies. Front Immunol 9, 2290, doi:10.3389/fimmu.2018.02290 (2018).

19 Cesarman, E. Gammaherpesviruses and lymphoproliferative disorders. Annual Review of Pathology: Mechanisms of Disease 9, 349–372 (2014).

20 Münz, C. Epstein Barr Virus Volume 2: One Herpes Virus: Many Diseases. Vol. 391 (2015).

21 Munz, C. Immune checkpoints in T cells during oncogenic gamma-herpesvirus infections. J Med Virol 95, e27840, doi:10.1002/jmv.27840 (2023).

22 Fisher, R. C. & Thorley-Lawson, D. A. Characterization of the Epstein-Barr virus-inducible gene encoding the human leukocyte adhesion and activation antigen BLAST-1 (CD48). Mol Cell Biol 11, 1614–1623, doi:10.1128/mcb.11.3.1614-1623.1991 (1991).

23 Thorley-Lawson, D. A., Ianelli, C., Klaman, L. D., Staunton, D. & Yokoyama, S. Function of CD48 and its regulation by Epstein-Barr virus. Biochem Soc Trans 21, 976–980, doi:10.1042/bst0210976 (1993).

24 Palendira, U. et al. Molecular pathogenesis of EBV susceptibility in XLP as revealed by analysis of female carriers with heterozygous expression of SAP. PLoS Biol 9, e1001187, doi:10.1371/journal.pbio.1001187 (2011).

25 Westhoff Smith, D., Chakravorty, A., Hayes, M., Hammerschmidt, W. & Sugden, B. The Epstein-Barr Virus Oncogene EBNA1 Suppresses Natural Killer Cell Responses and Apoptosis Early after Infection of Peripheral B Cells. mBio 12, e0224321, doi:10.1128/mBio.02243-21 (2021).

26 Green, M. & Michaels, M. G. Epstein-Barr virus infection and posttransplant lymphoproliferative disorder. Am J Transplant 13 Suppl 3, 41–54; quiz 54, doi:10.1111/ajt.12004 (2013).

27 El-Mallawany, N. K. & Rouce, R. H. EBV and post-transplant lymphoproliferative disorder: a complex relationship. Hematology Am Soc Hematol Educ Program 2024, 728–735, doi:10.1182/hematology.2024000583 (2024).

28 Campo, E. et al. The 2008 WHO classification of lymphoid neoplasms and beyond: evolving concepts and practical applications. Blood 117, 5019–5032, doi:10.1182/blood-2011-01-293050 (2011).

29 Jagadeesh, D., Woda, B. A., Draper, J. & Evens, A. M. Post Transplant Lymphoproliferative Disorders: Risk, Classification, and Therapeutic Recommendations. Current Treatment Options in Oncology 13, 122–136, doi:10.1007/s11864-011-0177-x (2012).

30 Swinnen, L. J. et al. Prospective study of sequential reduction in immunosuppression, interferon alpha-2B, and chemotherapy for posttransplantation lymphoproliferative disorder. Transplantation 86, 215–222, doi:10.1097/TP.0b013e3181761659 (2008).

31 Lanier, L. L. NK cell recognition. Annu Rev Immunol 23, 225–274, doi:10.1146/annurev.immunol.23.021704.115526 (2005).

32 Chen, S., Zhu, H. & Jounaidi, Y. Comprehensive snapshots of natural killer cells functions, signaling, molecular mechanisms and clinical utilization. Signal Transduct Target Ther 9, 302, doi:10.1038/s41392-024-02005-w (2024).

33 Vivier, E., Tomasello, E., Baratin, M., Walzer, T. & Ugolini, S. Functions of natural killer cells. Nat Immunol 9, 503–510, doi:10.1038/ni1582 (2008).

34 Orange, J. S. Formation and function of the lytic NK-cell immunological synapse. Nat Rev Immunol 8, 713–725, doi:10.1038/nri2381 (2008).

35 Bjorkstrom, N. K., Strunz, B. & Ljunggren, H. G. Natural killer cells in antiviral immunity. Nat Rev Immunol 22, 112–123, doi:10.1038/s41577-021-00558-3 (2022).

36 Wolf, N. K., Kissiov, D. U. & Raulet, D. H. Roles of natural killer cells in immunity to cancer, and applications to immunotherapy. Nat Rev Immunol 23, 90–105, doi:10.1038/s41577-022-00732-1 (2023).

37 Yodoi, J. et al. TCGF (IL 2)-receptor inducing factor(s). I. Regulation of IL 2 receptor on a natural killer-like cell line (YT cells). J Immunol 134, 1623-1630 (1985).

38 Cichocki, F. et al. Nicotinamide enhances natural killer cell function and yields remissions in patients with non-Hodgkin lymphoma. Sci Transl Med 15, eade3341, doi:10.1126/scitranslmed.ade3341 (2023).

39 Sanson, K. R. et al. Optimized libraries for CRISPR-Cas9 genetic screens with multiple modalities. Nat Commun 9, 5416, doi:10.1038/s41467-018-07901-8 (2018).

40 Doench, J. G. et al. Optimized sgRNA design to maximize activity and minimize off-target effects of CRISPR-Cas9. Nat Biotechnol 34, 184–191, doi:10.1038/nbt.3437 (2016).

41 Veillette, A. NK cell regulation by SLAM family receptors and SAP-related adapters. Immunol Rev 214, 22–34, doi:10.1111/j.1600-065X.2006.00453.x (2006).

42 McArdel, S. L., Terhorst, C. & Sharpe, A. H. Roles of CD48 in regulating immunity and tolerance. Clin Immunol 164, 10–20, doi:10.1016/j.clim.2016.01.008 (2016).

43 Wang, L. W. et al. Epstein-Barr-Virus-Induced One-Carbon Metabolism Drives B Cell Transformation. Cell Metab 30, 539–555 e511, doi:10.1016/j.cmet.2019.06.003 (2019).

44 Kinoshita, T. & Fujita, M. Biosynthesis of GPI-anchored proteins: special emphasis on GPI lipid remodeling. J Lipid Res 57, 6–24, doi:10.1194/jlr.R063313 (2016).

45 Staunton, D. E. et al. Blast-1 possesses a glycosyl-phosphatidylinositol (GPI) membrane anchor, is related to LFA-3 and OX-45, and maps to chromosome 1q21-23. J Exp Med 169, 1087–1099, doi:10.1084/jem.169.3.1087 (1989).

46 Seczynska, M. & Lehner, P. J. The sound of silence: mechanisms and implications of HUSH complex function. Trends in Genetics, 10.1016/j.tig.2022.12.005 (2023).

47 Seczynska, M., Bloor, S., Cuesta, S. M. & Lehner, P. J. Genome surveillance by HUSH-mediated silencing of intronless mobile elements. Nature 601, 440–445, doi:10.1038/s41586-021-04228-1 (2022).

48 Tchasovnikarova, I. A. et al. GENE SILENCING. Epigenetic silencing by the HUSH complex mediates position-effect variegation in human cells. Science 348, 1481–1485, doi:10.1126/science.aaa7227 (2015).

49 Li, D. Q. et al. MORC2 signaling integrates phosphorylation-dependent, ATPase-coupled chromatin remodeling during the DNA damage response. Cell Rep 2, 1657–1669, doi:10.1016/j.celrep.2012.11.018 (2012).

50 Tchasovnikarova, I. A. et al. Hyperactivation of HUSH complex function by Charcot-Marie-Tooth disease mutation in MORC2. Nat Genet 49, 1035–1044, doi:10.1038/ng.3878 (2017).

51 Ma, Y. et al. CRISPR/Cas9 Screens Reveal Epstein-Barr Virus-Transformed B Cell Host Dependency Factors. Cell Host Microbe 21, 580–591.e587, doi:10.1016/j.chom.2017.04.005 (2017).

52 Guo, R. et al. DNA methylation enzymes and PRC1 restrict B-cell Epstein-Barr virus oncoprotein expression. Nat Microbiol 5, 1051–1063, doi:10.1038/s41564-020-0724-y (2020).

53 Pross, H. F. & Jondal, M. Cytotoxic lymphocytes from normal donors. A functional marker of human non-T lymphocytes. Clin Exp Immunol 21, 226–235 (1975).

54 Goodman, J. J. The protection, preservation and strengthening of the abutment teeth through proper denture design. Part 2. *J Bergen Cty Dent Soc* 42, 7-13 (1975).

55 Montel, A. H., Morse, P. A. & Brahmi, Z. Upregulation of B7 molecules by the Epstein-Barr virus enhances susceptibility to lysis by a human NK-like cell line. Cell Immunol 160, 104–114, doi:10.1016/0008-8749(95)80015-b (1995).

56 Paul, S. & Lal, G. The Molecular Mechanism of Natural Killer Cells Function and Its Importance in Cancer Immunotherapy. Front Immunol 8, 1124, doi:10.3389/fimmu.2017.01124 (2017).

57 Lanier, L. L. Five decades of natural killer cell discovery. J Exp Med 221, doi:10.1084/jem.20231222 (2024).

58 Long, E. O., Kim, H. S., Liu, D., Peterson, M. E. & Rajagopalan, S. Controlling natural killer cell responses: integration of signals for activation and inhibition. Annu Rev Immunol 31, 227–258, doi:10.1146/annurev-immunol-020711-075005 (2013).

59 Kadri, N. et al. Dynamic Regulation of NK Cell Responsiveness. Curr Top Microbiol Immunol 395, 95–114, doi:10.1007/82_2015_485 (2016).

60 Dopkins, N. et al. A field guide to endogenous retrovirus regulatory networks. Molecular Cell 82, 3763–3768, 10.1016/j.molcel.2022.09.011 (2022).

61 Jensvold, Z. D., Flood, J. R., Christenson, A. E. & Lewis, P. W. Interplay between Two Paralogous Human Silencing Hub (HuSH) Complexes in Regulating LINE-1 Element Silencing. Nat Commun 15, 9492, doi:10.1038/s41467-024-53761-w (2024).

62 Timms, R. T., Tchasovnikarova, I. A. & Lehner, P. J. Position-effect variegation revisited: HUSHing up heterochromatin in human cells. Bioessays 38, 333–343, doi:10.1002/bies.201500184 (2016).

63 Meltzer, S. et al. gamma-Protocadherins control synapse formation and peripheral branching of touch sensory neurons. Neuron 111, 1776–1794 e1710, doi:10.1016/j.neuron.2023.03.012 (2023).

64 Hagelkruys, A. et al. The HUSH complex controls brain architecture and protocadherin fidelity. Sci Adv 8, eabo7247, doi:10.1126/sciadv.abo7247 (2022).

65 van Roy, F. Beyond E-cadherin: roles of other cadherin superfamily members in cancer. Nat Rev Cancer 14, 121–134, doi:10.1038/nrc3647 (2014).

66 Ito, M. et al. Killer cell lectin-like receptor G1 binds three members of the classical cadherin family to inhibit NK cell cytotoxicity. J Exp Med 203, 289–295, doi:10.1084/jem.20051986 (2006).

67 Nakamura, S. et al. Molecular basis for E-cadherin recognition by killer cell lectin-like receptor G1 (KLRG1). J Biol Chem 284, 27327–27335, doi:10.1074/jbc.M109.038802 (2009).

68 Consortium, E. P. An integrated encyclopedia of DNA elements in the human genome. Nature 489, 57–74, doi:10.1038/nature11247 (2012).

69 Luo, Y. et al. New developments on the Encyclopedia of DNA Elements (ENCODE) data portal. Nucleic Acids Res 48, D882–D889, doi:10.1093/nar/gkz1062 (2020).

70 Kagda, M. S. et al. Data navigation on the ENCODE portal. Nat Commun 16, 9592, doi:10.1038/s41467-025-64343-9 (2025).

71 Li, W. et al. Shifts in receptors during submergence of an encephalitic arbovirus. Nature 632, 614–621, doi:10.1038/s41586-024-07740-2 (2024).

72 Clark, L. E. et al. VLDLR and ApoER2 are receptors for multiple alphaviruses. Nature 602, 475–480, doi:10.1038/s41586-021-04326-0 (2022).

73 Wang, X., Xiong, H. & Ning, Z. Implications of NKG2A in immunity and immune-mediated diseases. Front Immunol 13, 960852, doi:10.3389/fimmu.2022.960852 (2022).

74 Lazetic, S., Chang, C., Houchins, J. P., Lanier, L. L. & Phillips, J. H. Human natural killer cell receptors involved in MHC class I recognition are disulfide-linked heterodimers of CD94 and NKG2 subunits. J Immunol 157, 4741–4745 (1996).

75 Katano, I. et al. Long-term maintenance of peripheral blood derived human NK cells in a novel human IL-15-transgenic NOG mouse. Sci Rep 7, 17230, doi:10.1038/s41598-017-17442-7 (2017).

76 Mitul, M. T. et al. A tonsil organoid model reveals Epstein-Barr virus infected germinal center B cell states during primary infection. bioRxiv, doi:10.64898/2026.01.15.699764 (2026).

77 Zhang, X. et al. Protocadherin gamma A3 is expressed in follicular lymphoma irrespective of BCL2 status and is associated with tumor cell growth. Mol Med Rep 14, 4622–4628, doi:10.3892/mmr.2016.5808 (2016).

78 Takata, K. et al. Duodenal follicular lymphoma: comprehensive gene expression analysis with insights into pathogenesis. Cancer Sci 105, 608–615, doi:10.1111/cas.12392 (2014).

79 Beckwith, M., Longo, D. L., O’Connell, C. D., Moratz, C. M. & Urba, W. J. Phorbol ester-induced, cell-cycle-specific, growth inhibition of human B-lymphoma cell lines. J Natl Cancer Inst 82, 501–509, doi:10.1093/jnci/82.6.501 (1990).

80 Spencley, A. L. et al. Co-transcriptional genome surveillance by HUSH is coupled to termination machinery. Mol Cell 83, 1623–1639 e1628, doi:10.1016/j.molcel.2023.04.014 (2023).

81 Lehner, P. J. Silencing by the HUSH Epigenetic Transcriptional Repressor Complex. Annu Rev Biochem 94, 361–386, doi:10.1146/annurev-biochem-020425-045352 (2025).

82 Seczynska, M. & Lehner, P. J. The sound of silence: mechanisms and implications of HUSH complex function. Trends Genet 39, 251–267, doi:10.1016/j.tig.2022.12.005 (2023).

83 Chougui, G. et al. HIV-2/SIV viral protein X counteracts HUSH repressor complex. Nat Microbiol 3, 891–897, doi:10.1038/s41564-018-0179-6 (2018).

84 Pedersen, S. F. et al. Inhibition of a Chromatin and Transcription Modulator, SLTM, Increases HIV-1 Reactivation Identified by a CRISPR Inhibition Screen. J Virol 96, e0057722, doi:10.1128/jvi.00577-22 (2022).

85 Yurkovetskiy, L. et al. Primate immunodeficiency virus proteins Vpx and Vpr counteract transcriptional repression of proviruses by the HUSH complex. Nat Microbiol 3, 1354–1361, doi:10.1038/s41564-018-0256-x (2018).

86 Martin, M. M. et al. Binding to DCAF1 distinguishes TASOR and SAMHD1 degradation by HIV-2 Vpx. PLoS Pathog 17, e1009609, doi:10.1371/journal.ppat.1009609 (2021).

87 Matkovic, R. et al. TASOR epigenetic repressor cooperates with a CNOT1 RNA degradation pathway to repress HIV. Nat Commun 13, 66, doi:10.1038/s41467-021-27650-5 (2022).

88 Prigozhin, D. M. et al. Periphilin self-association underpins epigenetic silencing by the HUSH complex. Nucleic Acids Res 48, 10313–10328, doi:10.1093/nar/gkaa785 (2020).

89 Zhu, Y., Wang, G. Z., Cingoz, O. & Goff, S. P. NP220 mediates silencing of unintegrated retroviral DNA. Nature 564, 278–282, doi:10.1038/s41586-018-0750-6 (2018).

90 Roubille, S. et al. The HUSH epigenetic repressor complex silences PML nuclear body-associated HSV-1 quiescent genomes. Proc Natl Acad Sci U S A 121, e2412258121, doi:10.1073/pnas.2412258121 (2024).

91 Meng, S. et al. ZNF638 represses the transcription of HBV closed circular DNA involving HUSH complex-mediated histone modifications of epigenetic silencing. Cell Commun Signal 24, doi:10.1186/s12964-026-02726-1 (2026).

92 Ngo, A. M. & Puschnik, A. S. Genome-Scale Analysis of Cellular Restriction Factors That Inhibit Transgene Expression from Adeno-Associated Virus Vectors. J Virol 97, e0194822, doi:10.1128/jvi.01948-22 (2023).

93 Das, A. et al. Epigenetic Silencing of Recombinant Adeno-associated Virus Genomes by NP220 and the HUSH Complex. J Virol 96, e0203921, doi:10.1128/JVI.02039-21 (2022).

94 Jayatissa, A. et al. The ERVK3-1 Microprotein Interacts with the HUSH Complex. Biochemistry 64, 3372–3381, doi:10.1021/acs.biochem.5c00023 (2025).

95 Bloor, S., Wit, N. & Lehner, P. J. RNA binding by Periphilin plays an essential role in initiating silencing by the HUSH complex. Nucleic Acids Res 53, doi:10.1093/nar/gkae1165 (2025).

96 Robbez-Masson, L. et al. The HUSH complex cooperates with TRIM28 to repress young retrotransposons and new genes. Genome Res 28, 836–845, doi:10.1101/gr.228171.117 (2018).

97 Braud, V. M. et al. HLA-E binds to natural killer cell receptors CD94/NKG2A, B and C. Nature 391, 795–799, doi:10.1038/35869 (1998).

98 Borrego, F., Ulbrecht, M., Weiss, E. H., Coligan, J. E. & Brooks, A. G. Recognition of human histocompatibility leukocyte antigen (HLA)-E complexed with HLA class I signal sequence-derived peptides by CD94/NKG2 confers protection from natural killer cell-mediated lysis. J Exp Med 187, 813–818, doi:10.1084/jem.187.5.813 (1998).

99 Lee, N. et al. HLA-E is a major ligand for the natural killer inhibitory receptor CD94/NKG2A. Proc Natl Acad Sci U S A 95, 5199–5204, doi:10.1073/pnas.95.9.5199 (1998).

100 Barrow, A. D., Martin, C. J. & Colonna, M. The Natural Cytotoxicity Receptors in Health and Disease. Front Immunol 10, 909, doi:10.3389/fimmu.2019.00909 (2019).

101 Brandt, C. S. et al. The B7 family member B7-H6 is a tumor cell ligand for the activating natural killer cell receptor NKp30 in humans. J Exp Med 206, 1495–1503, doi:10.1084/jem.20090681 (2009).

102 Jarahian, M. et al. Modulation of NKp30-and NKp46-mediated natural killer cell responses by poxviral hemagglutinin. PLoS Pathog 7, e1002195, doi:10.1371/journal.ppat.1002195 (2011).

103 Mavoungou, E., Held, J., Mewono, L. & Kremsner, P. G. A Duffy binding-like domain is involved in the NKp30-mediated recognition of Plasmodium falciparum-parasitized erythrocytes by natural killer cells. J Infect Dis 195, 1521–1531, doi:10.1086/515579 (2007).

104 Li, S. S. et al. The NK receptor NKp30 mediates direct fungal recognition and killing and is diminished in NK cells from HIV-infected patients. Cell Host Microbe 14, 387–397, doi:10.1016/j.chom.2013.09.007 (2013).

105 Lv, D. et al. The similar expression pattern of MHC class I molecules in human and mouse cerebellar cortex. Neurochem Res 39, 180–186, doi:10.1007/s11064-013-1204-z (2014).

106 Cebrian, C. et al. MHC-I expression renders catecholaminergic neurons susceptible to T-cell-mediated degeneration. Nat Commun 5, 3633, doi:10.1038/ncomms4633 (2014).

107 Muldoon, L. L. et al. Immunologic privilege in the central nervous system and the blood-brain barrier. J Cereb Blood Flow Metab 33, 13–21, doi:10.1038/jcbfm.2012.153 (2013).

108 Ning, Z., Liu, Y., Guo, D., Lin, W. J. & Tang, Y. Natural killer cells in the central nervous system. Cell Commun Signal 21, 341, doi:10.1186/s12964-023-01324-9 (2023).

109 Poli, A. et al. NK cells in central nervous system disorders. J Immunol 190, 5355–5362, doi:10.4049/jimmunol.1203401 (2013).

110 Wang, X. et al. Gamma protocadherins are required for survival of spinal interneurons. Neuron 36, 843–854, doi:10.1016/s0896-6273(02)01090-5 (2002).

111 Weiner, J. A., Wang, X., Tapia, J. C. & Sanes, J. R. Gamma protocadherins are required for synaptic development in the spinal cord. Proc Natl Acad Sci U S A 102, 8–14, doi:10.1073/pnas.0407931101 (2005).

112 Loh, K. H. et al. Proteomic Analysis of Unbounded Cellular Compartments: Synaptic Clefts. Cell 166, 1295–1307 e1221, doi:10.1016/j.cell.2016.07.041 (2016).

113 Phillips, G. R. et al. Gamma-protocadherins are targeted to subsets of synapses and intracellular organelles in neurons. J Neurosci 23, 5096–5104, doi:10.1523/JNEUROSCI.23-12-05096.2003 (2003).

114 Lin Chua, H. & Brahmi, Z. Expression of p58.2 or CD94/NKG2A inhibitory receptors in an NK-like cell line, YTINDY, leads to HLA Class I-mediated inhibition of cytotoxicity in the p58.2-but not the CD94/NKG2A-expressing transfectant. Cell Immunol 219, 57–70, doi:10.1016/s0008-8749(02)00578-6 (2002).

115 Tunbak, H. et al. The HUSH complex is a gatekeeper of type I interferon through epigenetic regulation of LINE-1s. Nat Commun 11, 5387, doi:10.1038/s41467-020-19170-5 (2020).

116 Billaud, M. et al. Low expression of lymphocyte function-associated antigen (LFA)-1 and LFA-3 adhesion molecules is a common trait in Burkitt’s lymphoma associated with and not associated with Epstein-Barr virus. Blood 75, 1827–1833 (1990).

117 Gregory, C. D., Murray, R. J., Edwards, C. F. & Rickinson, A. B. Downregulation of cell adhesion molecules LFA-3 and ICAM-1 in Epstein-Barr virus-positive Burkitt’s lymphoma underlies tumor cell escape from virus-specific T cell surveillance. J Exp Med 167, 1811–1824, doi:10.1084/jem.167.6.1811 (1988).

118 Mitra, B. et al. Characterization of target gene regulation by the two Epstein-Barr virus oncogene LMP1 domains essential for B-cell transformation. mBio 14, e0233823, doi:10.1128/mbio.02338-23 (2023).

119 Katano, H., Pesnicak, L. & Cohen, J. I. Simvastatin induces apoptosis of Epstein-Barr virus (EBV)-transformed lymphoblastoid cell lines and delays development of EBV lymphomas. Proc Natl Acad Sci U S A 101, 4960–4965, doi:10.1073/pnas.0305149101 (2004).

120 Park, J. H. & Faller, D. V. Epstein-Barr virus latent membrane protein-1 induction by histone deacetylase inhibitors mediates induction of intercellular adhesion molecule-1 expression and homotypic aggregation. Virology 303, 345–363, doi:10.1006/viro.2002.1638 (2002).

121 Wang, F. et al. Epstein-Barr virus latent membrane protein (LMP1) and nuclear proteins 2 and 3C are effectors of phenotypic changes in B lymphocytes: EBNA-2 and LMP1 cooperatively induce CD23. J Virol 64, 2309–2318, doi:10.1128/JVI.64.5.2309-2318.1990 (1990).

122 Ersing, I. et al. A Temporal Proteomic Map of Epstein-Barr Virus Lytic Replication in B Cells. Cell Rep 19, 1479–1493, doi:10.1016/j.celrep.2017.04.062 (2017).

123 Lacerda, J. F. et al. Human Epstein-Barr virus (EBV)-specific cytotoxic T lymphocytes home preferentially to and induce selective regressions of autologous EBV-induced B cell lymphoproliferations in xenografted C.B-17 scid/scid mice. J Exp Med 183, 1215-1228, doi:10.1084/jem.183.3.1215 (1996).

124 Moosmann, A. et al. B cells immortalized by a mini-Epstein-Barr virus encoding a foreign antigen efficiently reactivate specific cytotoxic T cells. Blood 100, 1755–1764 (2002).

125 Atallah-Yunes, S. A., Salman, O. & Robertson, M. J. Post-transplant lymphoproliferative disorder: Update on treatment and novel therapies. Br J Haematol 201, 383–395, doi:10.1111/bjh.18763 (2023).

126 Mahadeo, K. M. et al. Tabelecleucel for allogeneic haematopoietic stem-cell or solid organ transplant recipients with Epstein-Barr virus-positive post-transplant lymphoproliferative disease after failure of rituximab or rituximab and chemotherapy (ALLELE): a phase 3, multicentre, open-label trial. Lancet Oncol 25, 376–387, doi:10.1016/S1470-2045(23)00649-6 (2024).

127 Heslop, H. E. et al. Long-term outcome of EBV-specific T-cell infusions to prevent or treat EBV-related lymphoproliferative disease in transplant recipients. Blood 115, 925–935, doi:10.1182/blood-2009-08-239186 (2010).

128 Khanna, R. et al. Activation and adoptive transfer of Epstein-Barr virus-specific cytotoxic T cells in solid organ transplant patients with posttransplant lymphoproliferative disease. Proc Natl Acad Sci U S A 96, 10391–10396, doi:10.1073/pnas.96.18.10391 (1999).

129 Long, H. M. & Taylor, G. S. The T Cell Response to Epstein-Barr Virus. Curr Top Microbiol Immunol, doi:10.1007/82_2025_339 (2026).

130 Jabri, B. et al. TCR specificity dictates CD94/NKG2A expression by human CTL. Immunity 17, 487–499, doi:10.1016/s1074-7613(02)00427-2 (2002).

131 Andre, P. et al. Anti-NKG2A mAb Is a Checkpoint Inhibitor that Promotes Anti-tumor Immunity by Unleashing Both T and NK Cells. Cell 175, 1731–1743 e1713, doi:10.1016/j.cell.2018.10.014 (2018).

132 Chen, X. et al. Differential expression of inhibitory receptor NKG2A distinguishes disease-specific exhausted CD8(+) T cells. MedComm (2020) 3, e111, doi:10.1002/mco2.111 (2022).

133 Hislop, A. D., Taylor, G. S., Sauce, D. & Rickinson, A. B. Cellular responses to viral infection in humans: lessons from Epstein-Barr virus. Annu Rev Immunol 25, 587–617, doi:10.1146/annurev.immunol.25.022106.141553 (2007).

134 Pancho, A., Aerts, T., Mitsogiannis, M. D. & Seuntjens, E. Protocadherins at the Crossroad of Signaling Pathways. Front Mol Neurosci 13, 117, doi:10.3389/fnmol.2020.00117 (2020).

135 Higai, K., Imaizumi, Y., Suzuki, C., Azuma, Y. & Matsumoto, K. NKG2D and CD94 bind to heparin and sulfate-containing polysaccharides. Biochem Biophys Res Commun 386, 709–714, doi:10.1016/j.bbrc.2009.06.101 (2009).

136 Higai, K. et al. Binding affinities of NKG2D and CD94 to sialyl Lewis X-expressing N-glycans and heparin. Biol Pharm Bull 34, 8–12, doi:10.1248/bpb.34.8 (2011).

137 Muntasell, A. et al. Targeting NK-cell checkpoints for cancer immunotherapy. Curr Opin Immunol 45, 73–81, doi:10.1016/j.coi.2017.01.003 (2017).

138 Vietzen, H. et al. Inhibitory NKG2A(+) and absent activating NKG2C(+) NK cell responses are associated with the development of EBV(+) lymphomas. Front Immunol 14, 1183788, doi:10.3389/fimmu.2023.1183788 (2023).

139 Strati, P. et al. Off-the-shelf induced pluripotent stem-cell-derived natural killer-cell therapy in relapsed or refractory B-cell lymphoma: a multicentre, open-label, phase 1 study. Lancet Haematol 12, e505–e515, doi:10.1016/S2352-3026(25)00142-5 (2025).

140 Bachanova, V. et al. First-in-Human Phase I Study of Nicotinamide-Expanded Related Donor Natural Killer Cells for the Treatment of Relapsed/Refractory Non-Hodgkin Lymphoma and Multiple Myeloma. Biology of Blood and Marrow Transplantation 25, S175–S176, doi:10.1016/j.bbmt.2018.12.317 (2019).

141 Sun, Y., Chou, J., Dong, K. D., Gygi, S. P. & Gewurz, B. E. Insights into absence of lymphoma despite fulminant Epstein-Barr virus infection in patients with XIAP deficiency. JCI Insight 10, doi:10.1172/jci.insight.193787 (2025).

142 Gunesch, J. T. et al. CD56 regulates human NK cell cytotoxicity through Pyk2. Elife 9, doi:10.7554/eLife.57346 (2020).

143 Patro, R., Duggal, G., Love, M. I., Irizarry, R. A. & Kingsford, C. Salmon provides fast and bias-aware quantification of transcript expression. Nat Methods 14, 417–419, doi:10.1038/nmeth.4197 (2017).

144 Love, M. I., Huber, W. & Anders, S. Moderated estimation of fold change and dispersion for RNA-seq data with DESeq2. Genome Biol 15, 550, doi:10.1186/s13059-014-0550-8 (2014).

145 Sun, Y. et al. Epstein-barr virus latent membrane protein 1 targets cIAP1, cIAP2 and TRAF2 for proteasomal degradation to activate the non-canonical NF-kappaB pathway. PLoS Pathog 22, e1013898, doi:10.1371/journal.ppat.1013898 (2026).

