## Supplemental Figures and Tables for "The HUSH Complex Dictates EBV-transformed B cell Sensitivity to NK Cell Surveillance Through Repression of NKG2A Ligand γ-Proto-cadherin"

Figure S1

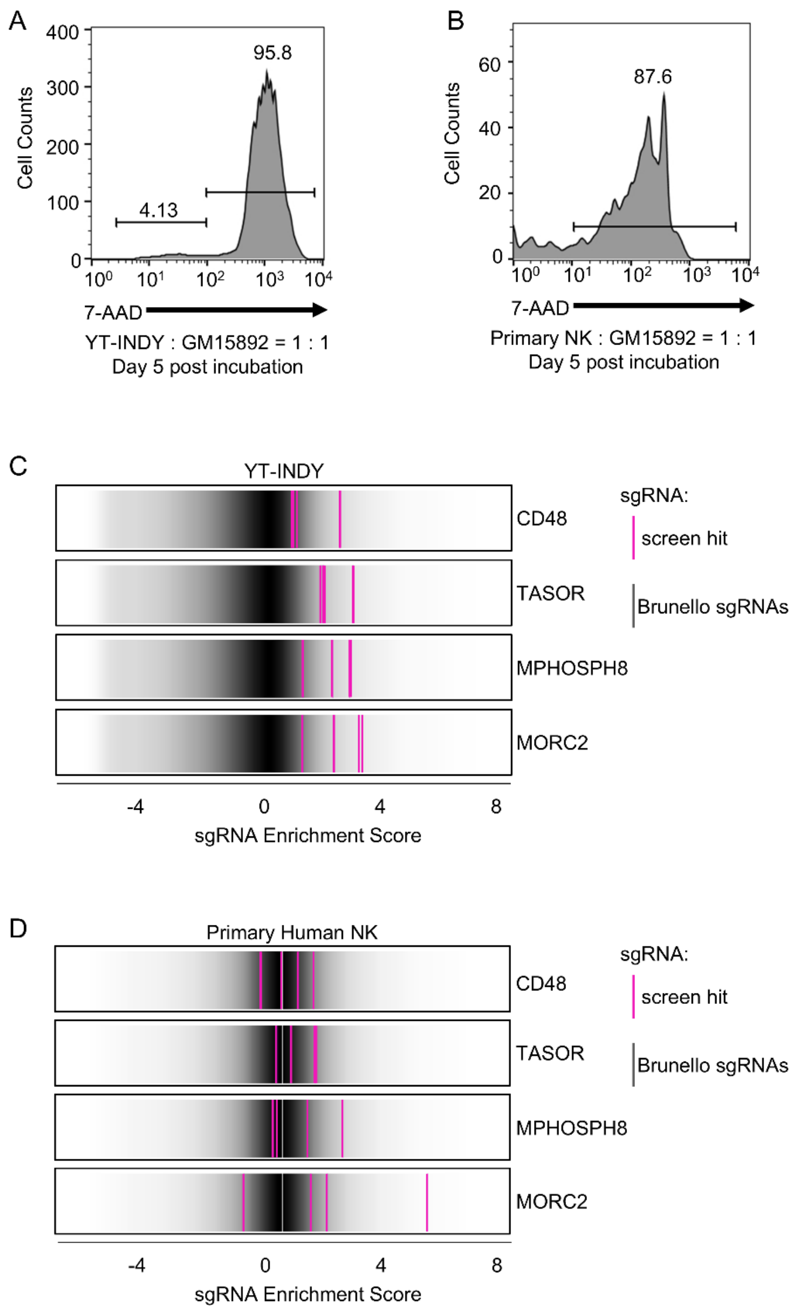


**Figure S1. CRISPR Screen for LCL Factors that Support NK Surveillance, Related to Figure 1.**

A) YT-INDY killing of GM15892 LCLs. FACS analysis of GM15892 cell vital dye 7-AAD uptake following five days of co-incubation with YT-INDY NK cells at an effector to target ratio of 1:1. Representative of n=3 replicates.

B) Primary human NK killing of GM15892 LCLs. FACS analysis of GM15892 cell vital dye 7-AAD uptake following five days of co-incubation with primary human peripheral blood IL-15/NAM treated NK cells at an effector to target ratio of 1:1. Representative of n=3 replicates.

C) Rug plot analysis of the indicated screen hit sgRNAs (pink) versus other Brunello library sgRNAs (gray) from the YT-INDY CRISPR screen.

D) Rug plot analysis of the indicated screen hit sgRNAs (pink) versus other Brunello library sgRNAs (gray) from the primary human NK CRISPR screen.

**Figure S2**

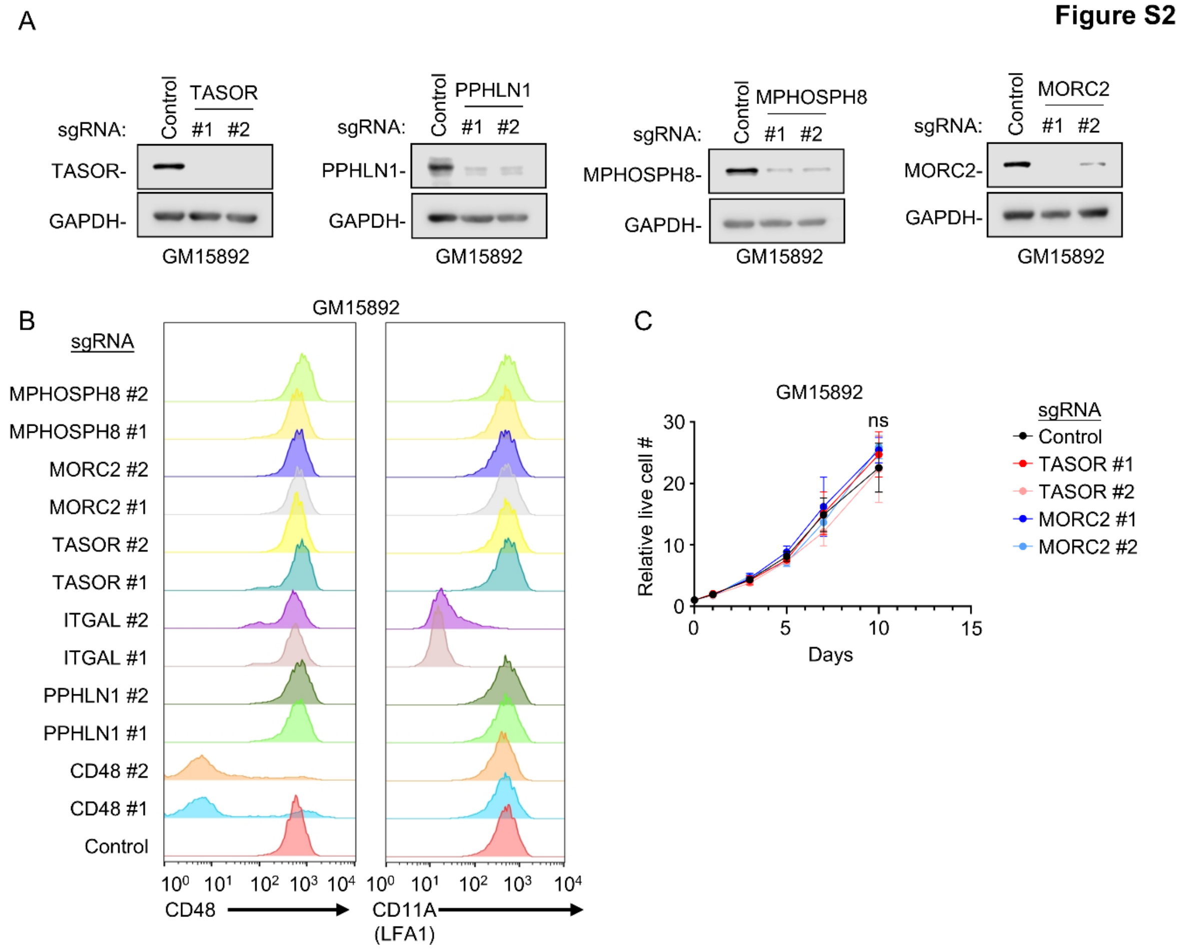


**Figure S2. Screen hit validation, related to Figure 1.**

A) Immunoblot analysis of whole cell lysates (WCL) from Cas9+ GM15892 LCLs that expressed screen the indicated control versus screen hit sgRNAs. Blots are representative of n=3 replicates.

B) FACS analysis of plasma membrane CD48 (left) versus ITGAL-encoded LFA-1 subunit CD11A (right) on Cas9+ GM15892 LCLs that expressed the indicated sgRNA. Data are representative of n=3 replicates.

C) Growth curve analysis of Cas9+ GM15892 cells that expressed the indicated sgRNAs. Shown are mean ± standard deviation (SD) live cell values from n=3 independent replicates.

Statistical significance was assessed by two-tailed unpaired Student's t test (C). ns, not significant.

**Figure S3**

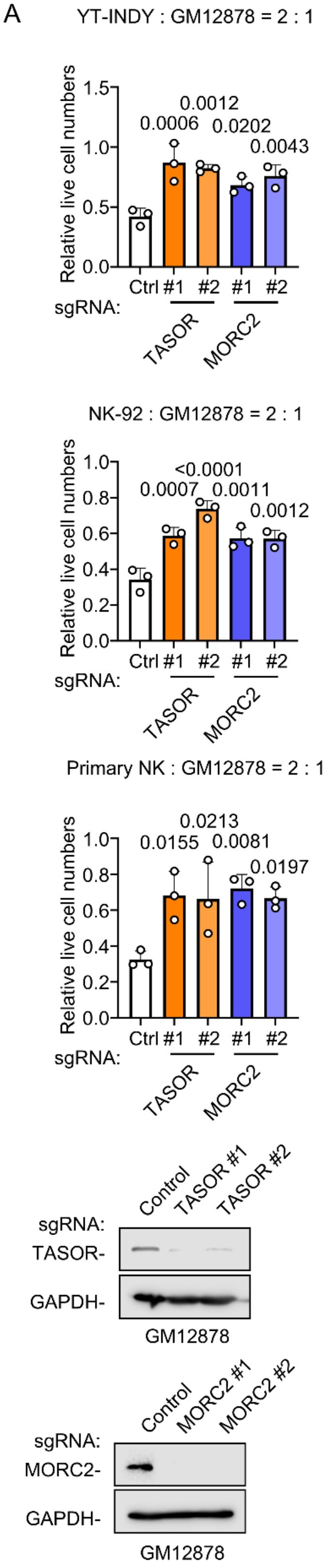
A
YT-INDY : GM12878 = 2 : 1
Relative live cell numbers
0.0006
0.0012
0.0202
0.0043
Ctrl #1 #2 #1 #2
sgRNA:
TASOR
MORC2
NK-92 : GM12878 = 2 : 1
0.0007
<0.0001
0.0011
0.0012
Primary NK : GM12878 = 2 : 1
0.0155
0.0213
0.0081
0.0197
Control
TASOR #1
TASOR #2
TASOR-
GAPDH-
GM12878
Control
MORC2 #1
MORC2 #2
MORC2-

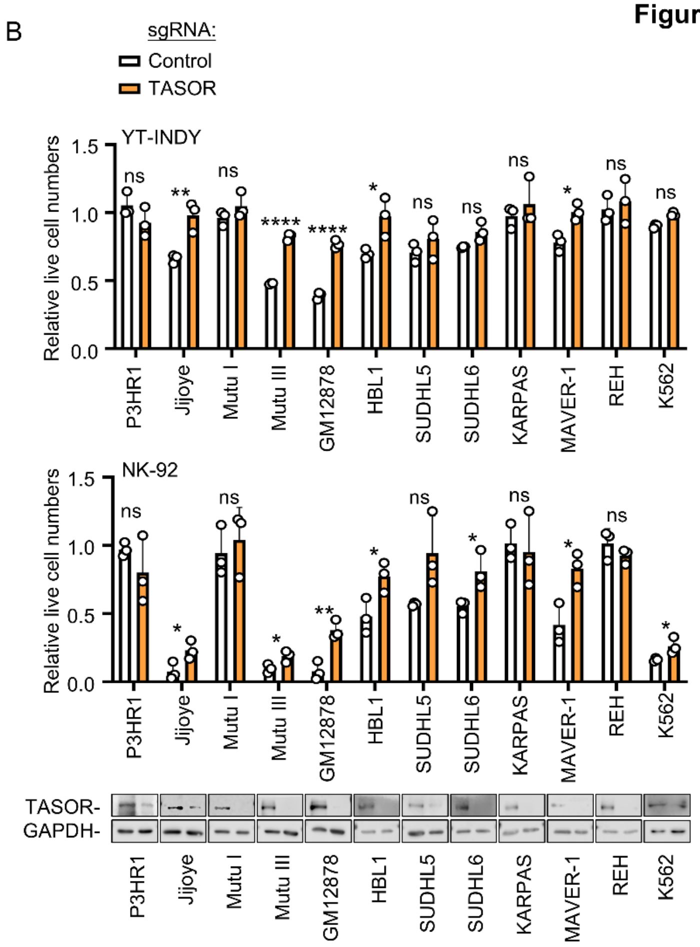
B
sgRNA:
Control
TASOR
YT-INDY
NK-92
Relative live cell numbers
P3HR1
Jijoye
Mutu I
Mutu III
GM12878
HBL1
SUDHL5
SUDHL6
KARPAS
MAVER-1
REH
K562
TASOR-
GAPDH-

**Figure S3. HUSH KO effects on target cell killing, related to Figure 2.**

A) HUSH KO effects on GM12878 LCL killing by NK. Mean + SD relative live cell numbers from n=3 replicates of Cas9+ GM12878 that expressed the indicated control (Ctrl), TASOR or MORC2 sgRNAs and co-cultured at a 2:1 ratio with YT-INDY (top), NK-92 (middle) or primary human IL-15/NAM treated NK (bottom) for 24 hours. Numbers were normalized to LCLs cultured in the absence of NK. Shown at bottom are representative immunoblots of WCL from GM12878 that expressed the indicated sgRNAs.

B) TASOR KO effects on B cell lymphoma or K562 killing by NK. Mean + SD relative live cell numbers from n=3 replicates of Cas9+ cells that expressed the indicated control (white bar) or TASOR (orange bar) sgRNAs and co-cultured at a 2:1 ratio with YT-INDY (top) or NK-92 (bottom) for 24 hours. Numbers were normalized to cells cultured in the absence of NK. Representative immunoblots of WCL from cells expressing control or TASOR sgRNAs are shown at bottom. Blots run separately are indicated by the borders.

Statistical significance was assessed by one-way ANOVA followed by Tukey’s multiple comparisons test (A) or two-tailed unpaired Student’s t test (B). ns, not significant, *P < 0.05, **P < 0.01, ***P < 0.001, ****P < 0.0001.

**Figure S4**

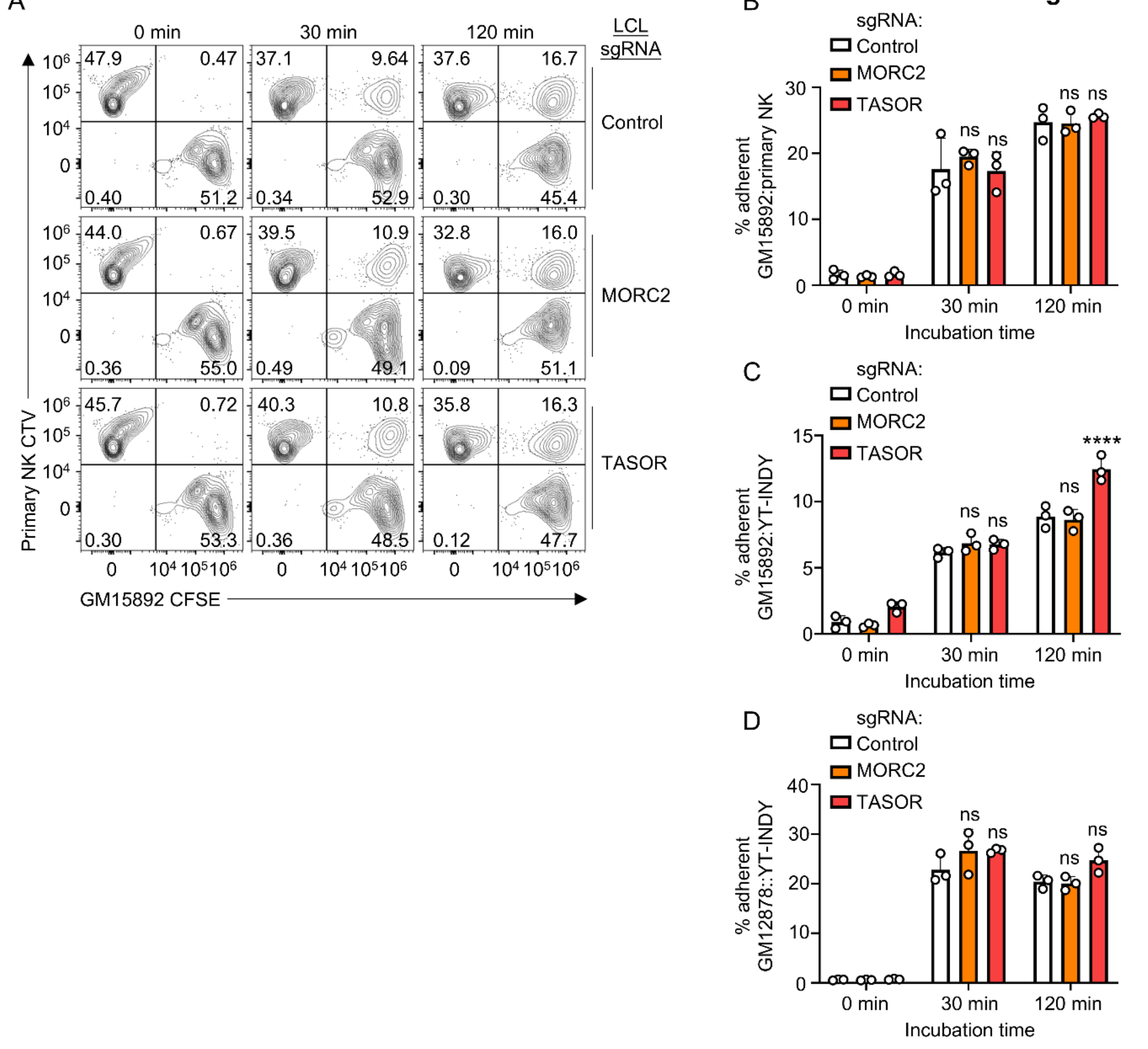


**Figure S4. LCL HUSH KO effects on NK adherence, Related to Figure 2.**

A) Effects of LCL HUSH NK on primary human NK adherence. Shown are representative FACS plots of *in vitro* adhesion assays, in which IL-15/NAM-activated primary human NK cells were co-cultured with Cas9+ GM15892 LCLs for the indicated minutes. NK were stained with CellTrace Violet (CTV) and GM15892 LCLs were stained with carboxyfluorescein succinimidyl ester (CFSE) just prior to the co-incubation.

B) Mean + SD percentages of adherent GM15892/primary human NK cell pairs from n=3 independent replicates as in (A).

C) Effects of LCL HUSH NK on YT-INDY GM15892 NK adherence. Shown are representative FACS plots of *in vitro* adhesion assays, in which YT-INDY NK cells were co-cultured with Cas9+ GM15892 LCLs for the indicated minutes. NK were stained with CTV and GM15892 LCLs were stained with CFSE just prior to the co-incubation. Mean + SD percentages of adherent GM15892/YT-INDY NK cell pairs from n=3 independent replicates.

D) Effects of LCL HUSH NK on GM12878 LCL YT-INDY NK adherence. Shown are representative FACS plots of *in vitro* adhesion assays, in which YT-INDY NK cells were co-cultured with Cas9+ GM12878 LCLs for the indicated minutes. NK were stained with CTV and GM12878 LCLs were stained with CFSE just prior to the co-incubation. Mean + SD percentages of adherent GM12878/YT-INDY NK cell pairs from n=3 independent replicates.

Statistical significance was assessed by one-way ANOVA followed by Tukey's multiple comparisons test (B-D). ns, not significant, *P < 0.05, **P < 0.01, ***P < 0.001, ****P < 0.0001.

Figure S5

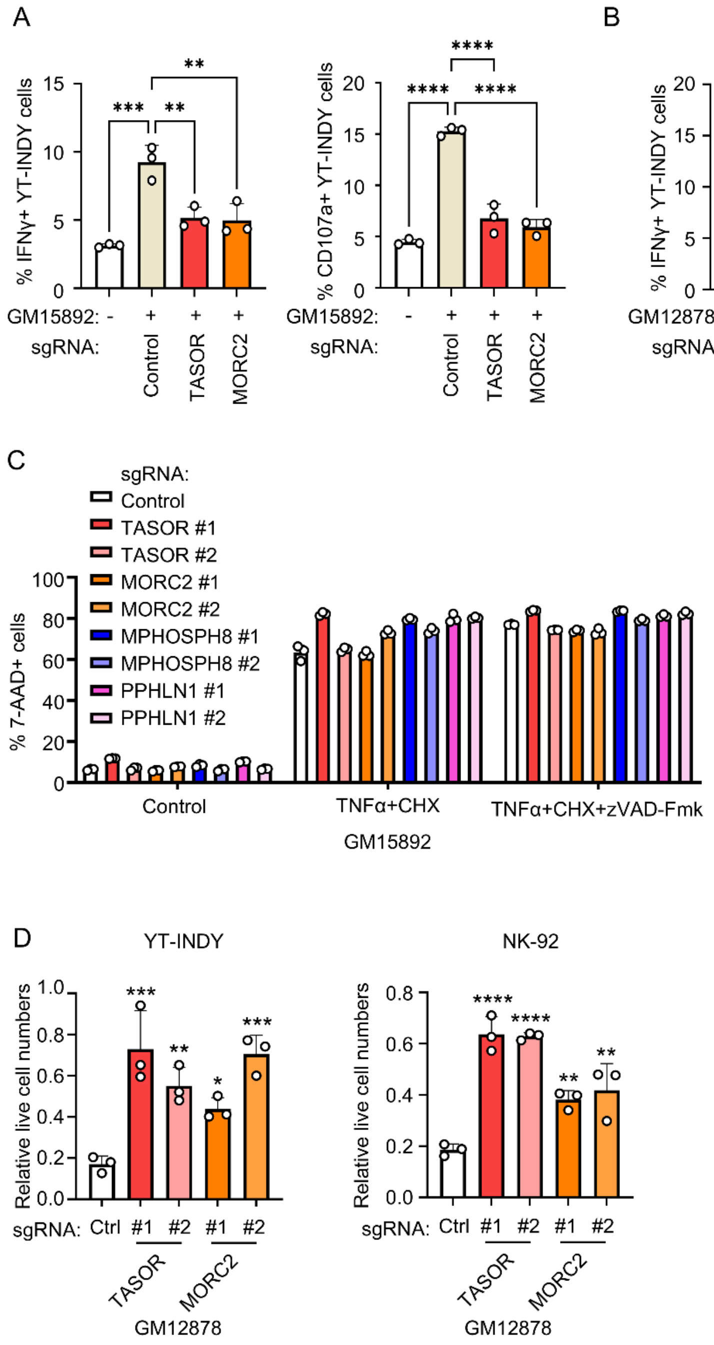


**Figure S5. HUSH KO effects on NK activation. Related to Figure 2.**

A) GM15892 LCL HUSH KO effects on YT-IDY IFNγ expression and plasma membrane CD107a. Mean + SD values from n=3 replicates of IFNγ+ (left) and CD107a+ (right) of YT-INDY cells that were cultured for 5 hours alone or with Cas9+ GM15892 LCLs that expressed the indicated sgRNAs. Cultures were treated with protein transport inhibitor for 5 hours prior to IFNγ analysis.

B) GM12878 LCL HUSH KO effects on YT-IDY IFNγ expression and plasma membrane CD107a. Mean + SD values from n=3 replicates of IFNγ+ (left) and CD107a+ (right) of YT-INDY cells that were cultured for 5 hours alone or with Cas9+ GM12878 LCLs that expressed the indicated sgRNAs. Cultures were treated with protein transport inhibitor for 5 hours prior to IFNγ analysis.

C) Analysis of HUSH KO effects on apoptosis and necroptosis induction. Shown are Mean + SD values of %7-AAD+ (dead) cells of Cas9+ GM15892 LCLs that expressed the indicated sgNRA and that were treated for 24 hours with TNFα (50ng/mL) and cycloheximide (CHX, 10μg/mL) for induction of apoptosis or with TNFα, CHX and caspase-inhibitor zVAD-Fmk (20μM) for induction of necroptosis.

D) Effects of exposure to HUSH KO LCL on subsequent NK killing. NK were pre-incubated with Cas9+ GM12878 with the indicated sgRNA for 4 days, and then used for killing assay with fresh control LCL targets. Mean + SD live cell numbers from n=3 replicates of GM12878 following co-culture with YT-INDY (left) or NK-92 (right) at a 2:1 E:T ratio for 24 hours. Values were normalized by live cell values of GM12878 cultured in the absence of NK.

Statistical significance was assessed by one-way ANOVA followed by Tukey's multiple comparisons test (A-D). ns, not significant, *P < 0.05, **P < 0.01, ***P < 0.001, ****P < 0.0001.

**Figure S6**

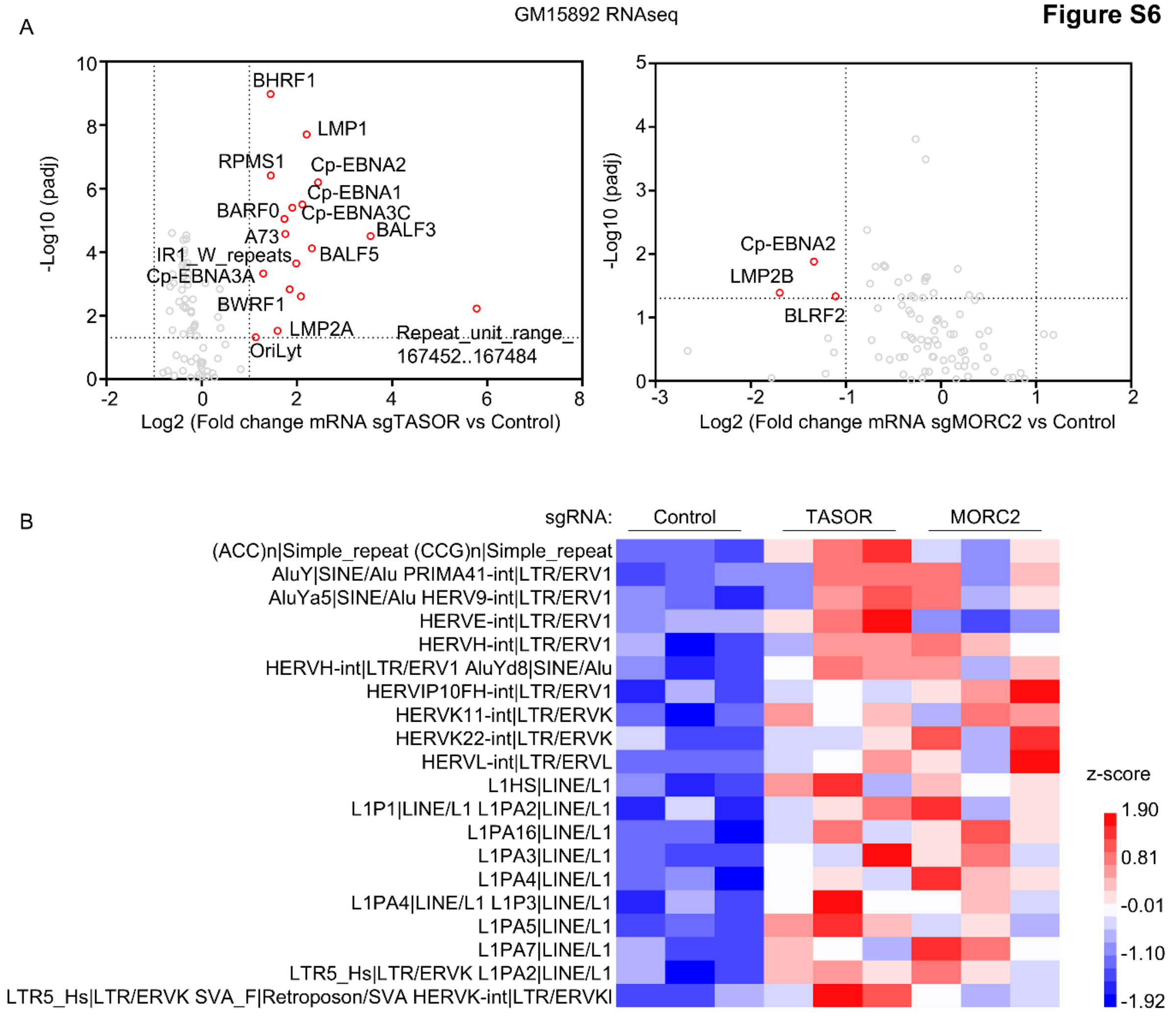


**Figure S6. HUSH KO effects on EBV and endogenous retroviral or LINE expression, related to Figure 3.**

A) Analysis of TASOR (left) or MORC2 (right) sgRNA effects on EBV gene expression. Shown are volcano plot analyses from n=3 RNAseq replicates of GM15892 that expressed control versus TASOR sgRNAs (left panel) or control versus MORC2 sgRNAs (right panel). Selected EBV genes are highlighted in red circles.

B) Heatmap analysis of the expression of human genome endogenous retroviral or LINE/L1 elements in GM15892 LCLs with control, TASOR or MORC2 sgRNAs from RNAseq analysis. Shown are the endogenous human genome elements whose expression was most highly changed by TASOR or MORC2 depletion.

**Figure S7**

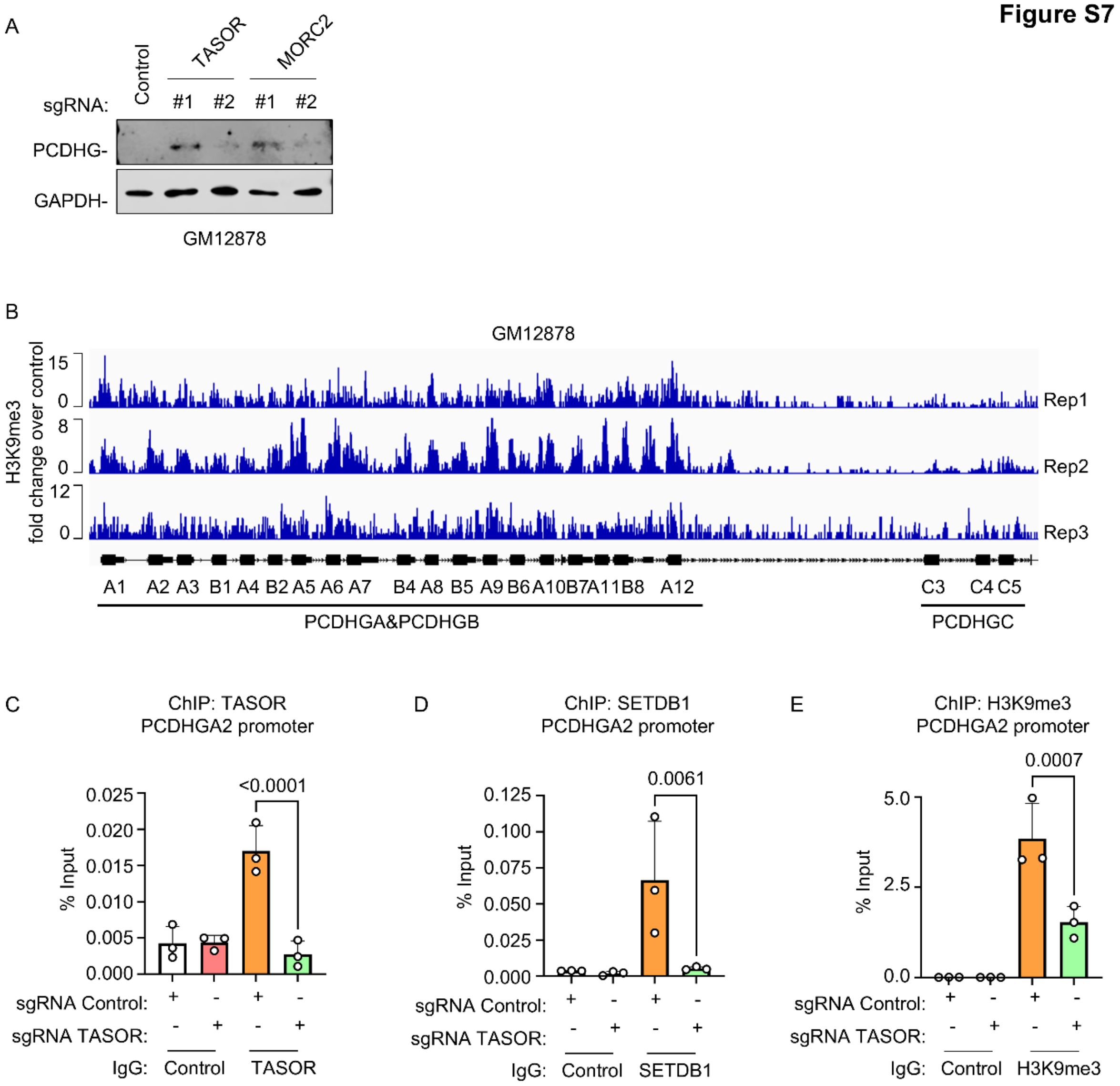


**Figure S7. Analysis of the LCL PCDHGA1 locus, related to Figure 3.**

A) Immunoblot analysis of WCL from GM12878 LCLs that expressed the indicated sgRNAs. Blots are representative of n=3 experiments.

B) H3K9me3 ChIP-seq tracks at the GM12878 LCL *PCDHG* locus. Data was obtained from the ENCODE project (accession: ENCSR000AOX).

C-E) ChIP-qPCR analysis of the GM15892 LCL *PCDHGA2* locus. Mean + SD values from n=3 ChIP-qPCR replicates of TASOR (C) or SETDB1 (D) occupancy or of H3K9me3 (E) levels at the *PCDHGA2* promoter region. Anti-TASOR, SETDB1 and H3K9me3 ChIP were performed on chromatin from GM15892 LCLs that expressed the indicated control or TASOR sgRNAs, followed by qPCR.

Statistical significance was assessed by two-tailed unpaired Student's t test (C-E).

**Figure S8**

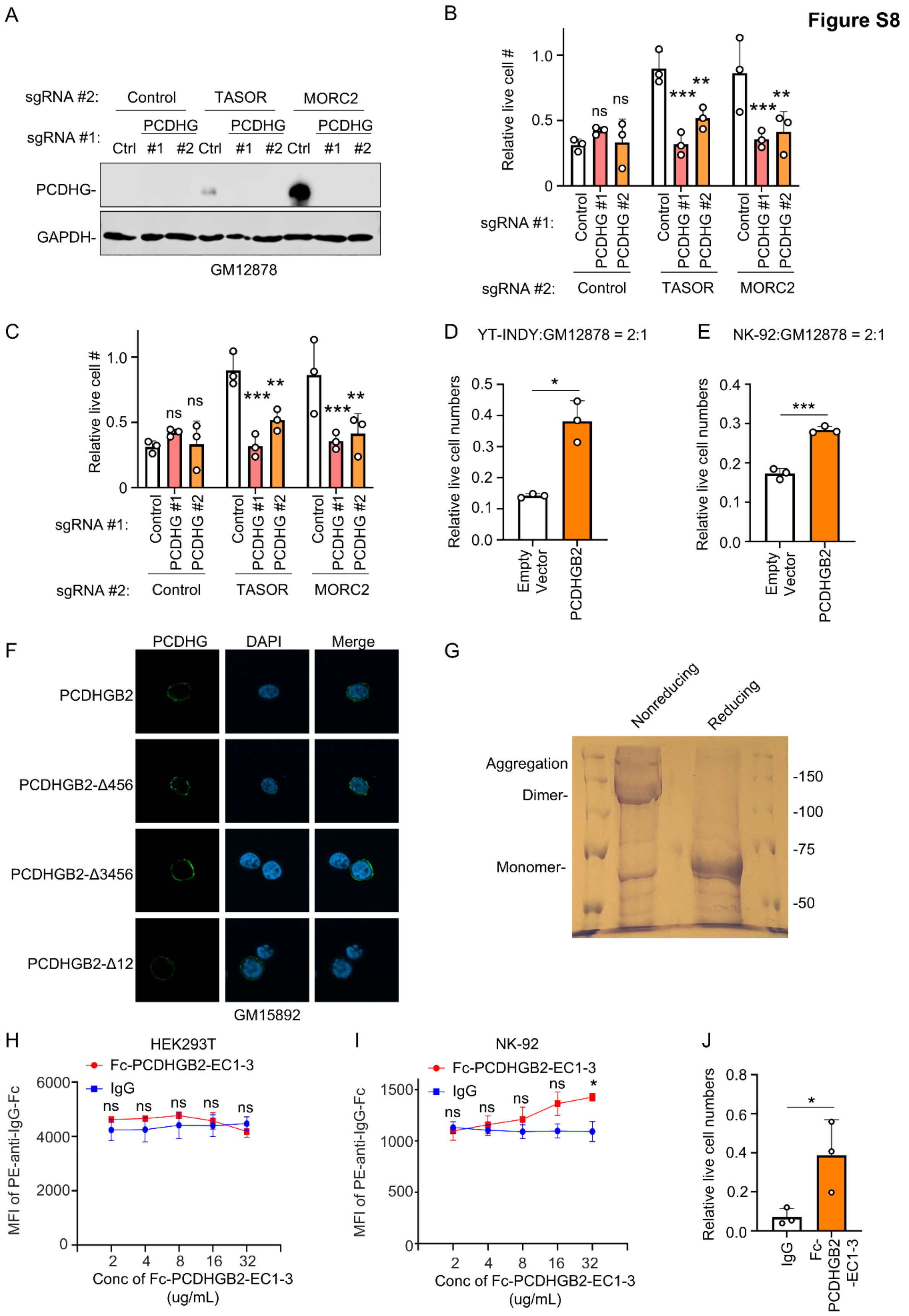
A
sgRNA #2: Control TASOR MORC2
sgRNA #1: PCDHG PCDHG PCDHG
Ctrl #1 #2 Ctrl #1 #2 Ctrl #1 #2
PCDHG-
GAPDH-
GM12878
B
Relative live cell #
1.0
0.5
0
ns
ns
***
**
***
**
sgRNA #1: Control PCDHG #1 PCDHG #2
sgRNA #2: Control TASOR MORC2
C
Relative live cell #
sgRNA #1:
sgRNA #2: Control TASOR MORC2
D YT-INDY:GM12878 = 2:1
Relative live cell numbers
Empty Vector
PCDHGB2
E NK-92:GM12878 = 2:1
F
PCDHG DAPI Merge
PCDHGB2
PCDHGB2-Δ456
PCDHGB2-Δ3456
PCDHGB2-Δ12
GM15892
G
Nonreducing Reducing
Aggregation
Dimer-
Monomer-
-150
-100
-75
-50
H HEK293T
Fc-PCDHGB2-EC1-3
IgG
MFI of PE-anti-IgG-Fc
Conc of Fc-PCDHGB2-EC1-3 (ug/mL)
I NK-92
J
IgG
Fc-PCDHGB2-EC1-3

**Figure S8. Analysis of PCDHG expression effects on NK-mediated LCL lysis, related to Fig. 4.**

A) Immunoblot analysis of WCL from GM12878 LCLs that expressed the indicated control or *PCDHG* common exon targeting sgRNAs, and that also expressed control, TASOR or MORC2 sgRNAs, as indicated.

B) Analysis of PCDHG knockout on HUSH KO resistance to YT-INDY mediated lysis. Mean + SD relative live cell values from n=3 replicates of Cas9+ GM12878 LCLs with the indicated sgRNAs following co-culture with YT-INDY at an E:T of 2:1 or culture alone for 24 hours.

C) Analysis of PCDHG knockout on HUSH KO resistance to NK92 mediated lysis. Mean + SD relative live cell values from n=3 replicates of Cas9+ GM12878 LCLs with the indicated sgRNAs following co-culture with NK-92 at an E:T of 2:1 or culture alone for 24 hours.

D) Analysis of PCDHGB2 expression effects on LCL protection from YT-INDY lysis. Mean + SD relative live cell numbers from n=3 replicates of GM12878 LCL that were electroporated with empty vector versus PCDGHB2 vectors and that were then co-cultured with YT-INDY at a 2:1 E:T ratio or alone for 24 hours. YT-INDY co-cultured LCL live cell values were normalized by values from identical LCLs cultured alone.

E) Analysis of PCDHGB2 expression effects on LCL protection from NK-92 lysis. Mean + SD relative live cell numbers from n=3 replicates of GM12878 LCL that were electroporated with empty vector versus PCDGHB2 vectors and that were then co-cultured with NK-92 at a 2:1 E:T ratio or alone for 24 hours. NK-92 co-cultured LCL live cell values were normalized by values from identical LCLs cultured alone.

F) Analysis of LCL PCDHGB2 transgene expression. Shown are representative confocal microscopy images of wildtype or truncation mutant PCDHGB2 cDNA expression in

GM15892 LCLs. Cells were transduced with lentiviral expression vectors and puromycin selected prior to immunostaining.

G) Analysis of recombinant Fc-PCDHGB2-EC1-3. Representative Coomassie blue stained gel loaded with purified Fc-PCDHGB2-EC1-3 loaded in non-reducing versus reducing loading buffer with 5% β-mercaptoethanol.

H) Analysis of Fc-PCDHGB2-EC1-3 labeling of HEK-293 plasma membranes. Shown are Mean ± SD MFI values from n=3 replicates of FACS analysis of control IgG versus Fc-PCDHGB2-EC1-3 stained HEK-293. Cells were treated with the indicated IgG versus Fc-PCDHGB2-EC1-3 concentrations and then stained with PE-tagged anti-IgG Fc prior to FACS analysis.

I) Analysis of Fc-PCDHGB2-EC1-3 labeling of NK-92 plasma membranes. Shown are Mean ± SD MFI values from n=3 replicates of FACS analysis of control IgG versus Fc-PCDHGB2-EC1-3 stained NK-92 cells. Cells were pretreated with Fc receptor blocking solution for 15 minutes, then treated with the indicated IgG versus Fc-PCDHGB2-EC1-3 concentrations and stained with PE-tagged anti-IgG Fc prior to FACS analysis.

J) Analysis of Fc-PCDHGB2-EC1-3 effects on NK-92 mediated LCL killing. Mean + SD relative live cell numbers from n=3 replicates of GM15892 cultured alone or with NK-92 that were pre-treated with 16 IgG or with Fc-PCDHGB2-EC1-3 μg/mL for 24 hours. LCLs and NK-92 were co-cultured at a 2:1 E:T ratio for 24 hours.

Statistical significance was assessed by two-tailed unpaired Student's t test (D,E,J) or two-way ANOVA followed by Tukey's multiple comparisons test (B,C,H,I). *P < 0.05, **P < 0.01, ***P<0.001, ****P < 0.0001, ns, not significant.

**Figure S9**

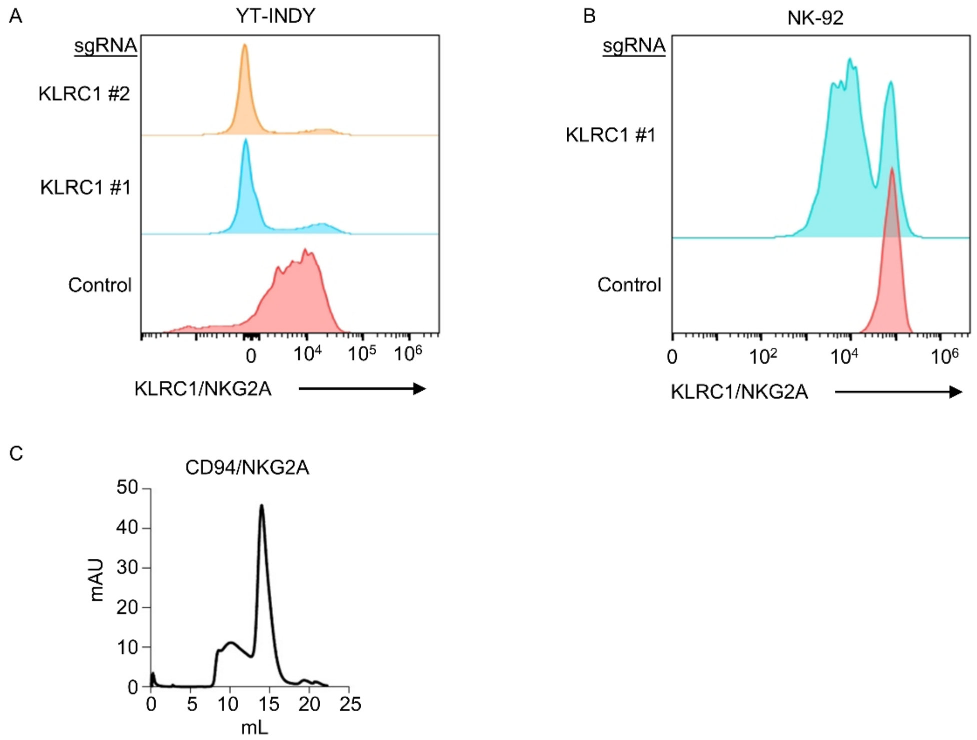


**Figure S9. Analysis of CRISPR depletion of KLRC1/NKG2A, related to Figure 5.**

A) Representative FACS analysis of plasma membrane NKG2A expression on Cas9+ YT-INDY cells that expressed control or independent screen hit KLRC1 sgRNAs.

B) Representative FACS analysis of plasma membrane NKG2A expression on Cas9+ NK-92 cells that expressed control or KLRC1 sgRNAs.

C) Size-exclusion chromatogram (SEC) elution traces of recombinant CD94/NKG2A.

Figure S10

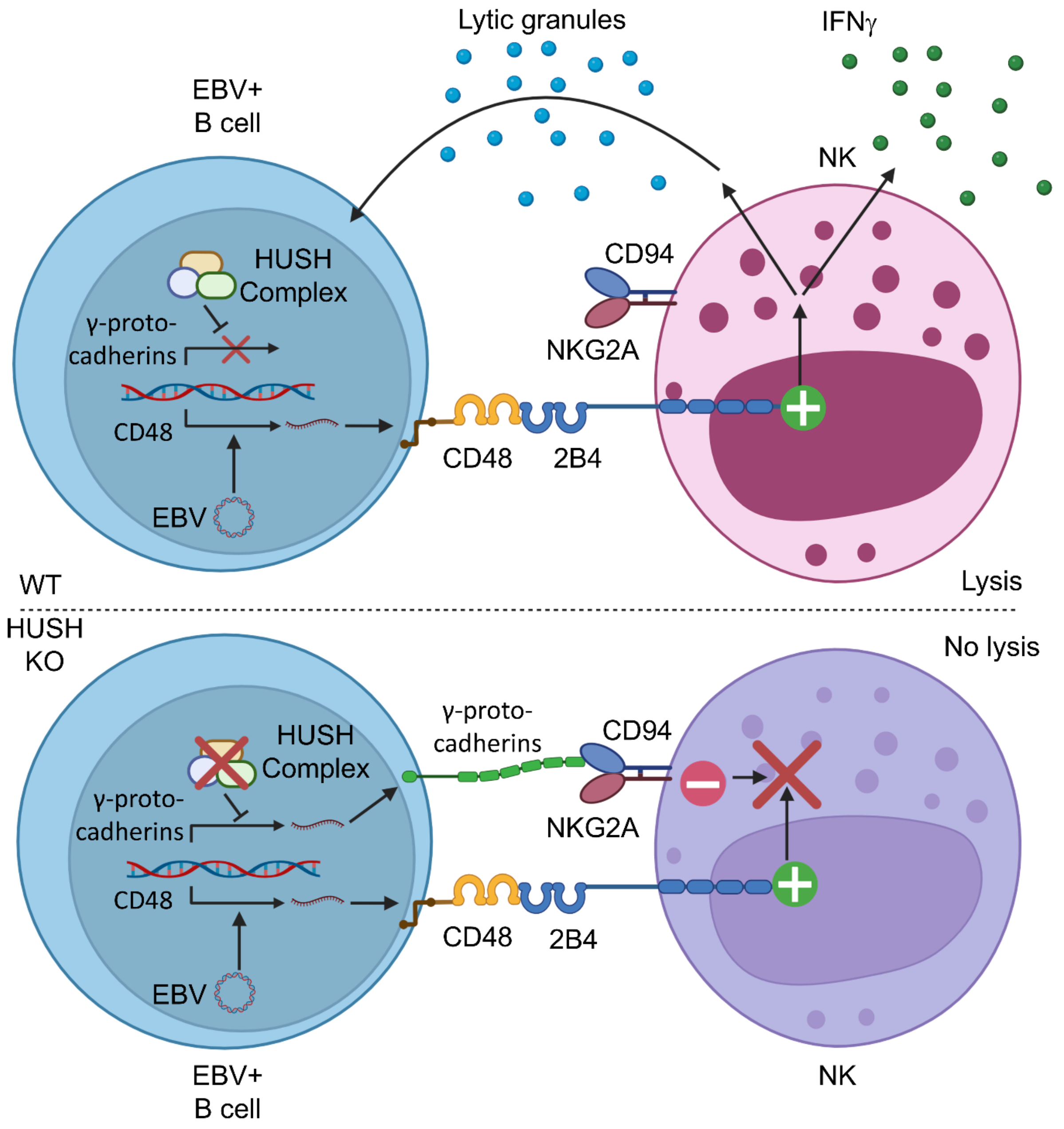


**Figure S10. Schematic model of key HUSH roles in PCDHG expression and in EBV-transformed lymphoblastoid B cell NK surveillance.**

EBV transformed B cells highly express CD48, which stimulates NK cell activation and infected B-cell lysis. The HUSH complex represses PCDHG expression to support EBV-transformed B cell driven NK activation. In the absence of HUSH activity, PCDHG is de-repressed, traffics to the plasma membrane and associates with NKG2A/CD94 to deliver an inhibitory signal that overcomes CD48/2B4 NK activation. This serves to support infected B cell escape from NK surveillance.

**Table S1. sgRNA or crRNA sequences used in this study**

| sgRNA name | sgRNA sequence |
|---|---|
| *CD48* #1 | CTGGTCGAAAGTATAAAACC |
| *CD48* #2 | TCACTTGGTACATATGACCG |
| *TASOR* #1 | TCTGTCTATTTATTACCCGG |
| *TASOR* #2 | ATAAATATAACTATACCTTG |
| *PPHLN1* #1 | AGGTGTTAGACAAACCCAGT |
| *PPHLN1* #2 | CTTCCCATTATGCGAGAGAG |
| *MPHOSPH8* #1 | CATAGACGATCACAAAACCA |
| *MPHOSPH8* #2 | GTCCACGGACTCAGCCGAGG |
| *MORC2* #1 | ACATTAGAAGTACGCCTAGG |
| *MORC2* #2 | GTGAGCTGCGTTACAAACGC |
| *ITGAL* #1 | TCTGGCGGAAGAGGTAACAC |
| *ITGAL* #2 | GCCACCGGACCAGAAGACGG |
| *PCDHG* #1 | ACTGGCGGACGCCAAGATCA |
| *PCDHG* #2 | TGCTGATGGGAGCTCCACCC |
| *KLRC1* #1 | TGAACAGGAAATAACCTATG |
| *KLRC1* #2 | GACAAAACCTATCACTGCAA |
| crRNA name | crRNA sequence |
| Control | CGTTAATCGCGTATAATACG |
| *MORC2* #1 | ACCACTCACGAATTCTTGTT |
| *MORC2* #2 | CCACCTGGAATGCTCGGACC |

**Table S2. ChIP-qPCR primer sequences used in this study**

| sgRNA name | sgRNA sequence |
|---|---|
| *PCDHGA1* FWD | GGAATGCAGTAACTGGTTAGGA |
| *PCDHGA1* REV | CCTGTGTGTACTGCCTGTATC |

| | |
|---|---|
| *PCDHGA2* FWD | CGCCTTGAAAGTCAACAAAGG |
| *PCDHGA2* REV | GACGCTCAGAGGCCTAAATAAT |
